# Lack of co-ordination of stomatal, hydraulic and leaf browning traits in 16 perennial Australian grass species of differing climate origins

**DOI:** 10.64898/2026.07.04.736528

**Authors:** Arjunan Krishnananthaselvan, Vinod Jacob, Jinyan Yang, Brendan Choat, Elise Pendall, Sally A. Power, David T. Tissue, Belinda E. Medlyn

## Abstract

Grasslands are vulnerable to increasing drought with global warming, but process-based models lack the mechanistic knowledge required to predict the magnitude of drought impacts. While a plant hydraulics framework has been successful in advancing process understanding of drought responses in trees, and how drought responses vary across rainfall gradients, similar approaches have rarely been applied to grasses.

Here, we quantified the progression of key drought response processes in sixteen dominant perennial grasses (seven C_3_ and nine C_4_) with differing climatic origins across eastern Australia.

We found that stomatal closure, hydraulic impairment and leaf browning occurred concurrently, in contrast to the progressive sequence typically observed in trees. We also found that drought response traits were not correlated with species’ climate of origin.

The early impairment of leaf hydraulic conductance and leaf browning along with the lack of correlation with climate of origin suggest that grasses may employ fundamentally different strategies to adapt to low water availability than trees. These results highlight the need for grass-specific parameterization of drought responses in process-based models.

## Introduction

Climate change is predicted to increase the frequency and severity of drought events (Dai, 2013; Masson-Delmotte et al., 2021), which is likely to have significant impacts on terrestrial vegetation structure and function. Grassland ecosystem productivity is highly sensitive to changes in precipitation (Huxman et al., 2004; Knapp et al., 2015, 2017; Wilcox et al., 2017), but predicting the response of grasslands to drought with vegetation models has proved challenging (De Kauwe et al., 2017; Gerten et al., 2008; Paschalis et al., 2020). One major obstacle to model development is the lack of data on the underlying physiological processes driving drought responses. Significant advances have been made in understanding and predicting responses of tree species to drought in recent years through investigation of plant hydraulics (Bartlett et al., 2016; Creek et al., 2018; Gauthey et al., 2020; Peters et al., 2021; Skelton et al., 2015; Trueba et al., 2019), which has enabled the development of new mechanistic models to describe drought impacts (Cochard et al., 2021; Sabot et al., 2020; Sperry et al., 2017; Venturas et al., 2018). However, there has been considerably less attention paid to grass species, and consequently fewer advances in mechanistic modelling. Our goal in this paper was to measure drought response parameters of key plant processes in a range of perennial grass species occurring across an aridity gradient, and test to what extent the patterns in these parameters differ from those observed in trees.

Perennial grasses are dominant in most sub-humid to semi-arid grasslands, including those in Australia. Like trees, perennial grasses need adaptations to survive extended periods of low moisture availability. However, the ways in which they adapt may be different. Although their physiological function is similar, perennial grasses differ strongly from trees in their morphological and life history characteristics (Donzelli et al., 2013; VanderWeide et al., 2014). The shallower root systems of grasses limit their ability to access deep water reserves, making them dependent on the more variable topsoil moisture (Kulmatiski et al., 2010; Ludwig et al., 2004). The construction cost for the hydraulic pathway of trees is much higher than that of grasses, and the loss of xylem conductivity in trees is correspondingly costly (Choat et al., 2018; Johnson et al., 2018), requiring significant investment to rebuild (Gauthey et al., 2022). In contrast, foliar senescence is an effective drought escape strategy in perennial grasses in seasonally dry grasslands, since they can re-sprout in the subsequent wet season (Blumenthal et al., 2021; T. W. Ocheltree et al., 2020; VanderWeide et al., 2014). Given these differences in morphology and life history, it may be expected that relationships among drought response traits would differ between perennial grasses and trees.

The key mechanisms involved in plant drought responses are stomatal closure, impairment of hydraulic conductance and leaf loss (Choat et al., 2018; McDowell et al., 2022). The physiological traits that we focus on here are the threshold water potential values commonly used to describe the progression of these processes with water stress: for example, P_gs50_ denotes the leaf water potential inducing 50% loss of stomatal conductance. In woody species, these processes have been shown to follow a consistent temporal sequence in which stomatal closure commences first, followed by hydraulic impairment (Bartlett et al., 2016; Trueba et al., 2019). The threshold of leaf water potential for stomatal closure (P_gs50_) occurs above the threshold for loss of leaf hydraulic conductivity (P_kl50_) which in turn occurs in advance of that for stem xylem embolism (P_50_). Stomatal closure thus occurs before xylem embolism, thereby leading to a positive hydraulic safety margin (HSM), defined as (P_gs88_ – P_50_) (Blackman, Creek, et al., 2019; Creek et al., 2018; Li et al., 2018, 2020; Skelton et al., 2015). This sequence of events is thought to minimize the risk of loss of hydraulic conductivity. This coordination further extends to the leaf turgor loss point (TLP), which represents the capacity of a leaf to maintain cell turgor pressure during desiccation. TLP is recognized as a key driver of stomatal downregulation (Salleo et al., 2000; Bartlett et al., 2012, 2014), typically occurring before significant stomatal closure (Blackman et al., 2010; Bourne et al., 2017; Farrell et al., 2017; Li et al., 2018; Yao et al., 2021).

Finally, leaf shedding may occur following the onset of xylem embolism in evergreen woody species (Blackman *et al*., 2019). Leaf shedding during extreme drought benefits the plant in two ways, through avoidance of respiratory loss of stored carbohydrates, and avoidance of transpirational loss of water (Vico et al., 2017). Typically, in trees, leaf shedding commences after the onset of xylem embolism; for example, Blackman et al. (2019) found that the water potential for the onset of leaf shedding corresponded to P_50_ across a suite of eucalypt species from different climate origins. In grasses, leaves brown off, rather than being shed, as drought progresses (T. W. Ocheltree et al., 2020; VanderWeide et al., 2014). In this study, we characterise this process using P_Brx_, defined as the leaf water potential at x% loss of canopy greenness, and tested when leaf browning occurred in the sequence of drought responses.

It is widely observed that the above traits vary in concert in woody species according to their climate of origin, particularly mean annual precipitation (MAP) and aridity index (AI) (Blackman et al., 2014; Gleason et al., 2016; Li et al., 2018; Nardini & Luglio, 2014; Peters et al., 2021). In both regional and global scale studies, tree species originating from dry environments tend to have lower P_50_ values, indicating greater tolerance for water stress, than species originating from wet environments (Choat et al., 2012; Li et al., 2018; Nardini & Luglio, 2014; Peters et al., 2021; Skelton et al., 2021). Stomatal closure thresholds and turgor loss point also typically co-vary with P_50_ across an aridity gradient (Blackman et al., 2012; Li et al., 2018).

In contrast, for grasses, the sequence of drought responses is less well characterized. It is not clear whether grasses typically maintain a positive or a negative HSM. However, there is some evidence for a positive HSM; e.g., in one major study of nine C_4_ grass species from Kansas, USA, Ocheltree *et al*. (2016) found that P_gs50_ was higher than P_kl50_ in all species, indicating a positive HSM. In contrast, Holloway-Phillips & Brodribb. (2011) measured drought-induced stomatal closure and hydraulic impairment in four cultivars of two forage grass species, *Lolium multiflorum* and *Festuca arundinacea*, and reported stomatal closure occurring well after the onset of hydraulic impairment in three out of four cultivars. However, both studies used the rehydration kinetics approach to measure loss of hydraulic conductivity in leaves. This method measures the total loss of hydraulic conductivity from both xylary and extra-xylary pathways, where the extra-xylary pathways may be more dynamic than those involving xylem embolism. Jacob et al. (2022) used an optical technique to monitor xylem embolism with increasing drought stress in five perennial pasture grasses and found that stomatal closure occurred before the onset of xylem embolism in all species. Thus, collectively, measured declines in leaf hydraulic conductance in grass species may reflect extra-xylary limitations rather than xylem cavitation, particularly when assessed using rehydration kinetics.

There is also relatively little information available on the relationship between hydraulic traits and climate of origin in grasses. Craine et al. (2013), in a survey of 426 species, found a weak relationship between P_gs50_ and mean annual precipitation (MAP) of origin. Ocheltree et al. (2016) did not find any relationship for either TLP or P_50_ and MAP of origin among nine C_4_ grass species in the USA. Baird et al. (2025) found that C_3_ grasses native to drier climates have higher rates of photosynthesis, stomatal conductance and leaf hydraulic conductance under standard well-watered conditions, but did not test drought response traits. Hence, more extensive studies are needed to establish general patterns.

Comparative studies on grasses need to consider differences related to photosynthetic pathway. Typically, C_3_ grasses exhibit higher operating stomatal conductance than C_4_ grasses (Taylor et al., 2011), and their drought responses may also differ. For example, Ripley et al. (2010) and Taylor et al. (2011) found a more rapid reduction in *g*_s_ with drought in C_3_ than in C_4_ grasses. Taylor et al. (2011) also found that C_4_ grasses exhibited higher leaf mortality during drought than C_3_ grasses. Drought sensitivity may thus vary between C_3_ and C_4_ species; however, it remains unclear whether drought response traits, and their sequence of occurrence, differ systematically with photosynthetic pathway.

A further component of trait co-ordination is the trade-off between hydraulic efficiency and safety, which has been used to explain the potential susceptibility of tree species to hydraulic failure. In tree stems, there is a trade-off, both within and across species, between the capacity of stem vessels to supply water to leaves (hydraulic efficiency) and their resistance to hydraulic failure (hydraulic safety) (Gleason et al., 2016; Jacobsen et al., 2007; Li et al., 2018). The existence of this trade-off in leaves is still under debate (Blackman et al., 2010; Nardini et al., 2012), and there is relatively little information available for grasses. Ocheltree et al. (2016) identified the existence of a safety vs. efficiency trade-off in leaves of nine C_4_ grass species, showing that across species, high hydraulic conductance was associated with high embolism vulnerability. However, this trade-off is not always observed (e.g. Jacob et al. 2022) and it is not yet possible to draw general conclusions on the hydraulic efficiency vs. safety trade-off in grasses.

Given this lack of data on the physiological drought response mechanisms in grasses, we aimed to quantify key drought response parameters in a suite of C_3_ and C_4_ grasses occurring over a broad rainfall gradient in eastern Australia. We aimed to answer the following questions: (1) In what order do the thresholds for stomatal closure, hydraulic failure and leaf browning occur in perennial grasses, and does this sequence differ from that typically observed in trees? (2) Do the traits characterizing the progression of stomatal closure, hydraulic impairment and leaf browning differ systematically between C_3_ and C_4_ grasses? (3) Do these traits vary with climate of origin across a gradient of rainfall or aridity? and (4) Do hydraulic safety and efficiency trade-offs exist in these grasses?

## Materials and Methods

Stomatal closure, hydraulic impairment and leaf browning drought response traits of 16 common grass species in Australia (Table 1) were quantified. Grasses were grown inside a polytunnel facility at Hawkesbury Institute for the Environment, Western Sydney University (33° 33′ S, 150° 44′ E, Richmond, NSW, Australia) under well-watered conditions for a minimum of three months, after which drought was imposed by withholding irrigation.

**Table 1.** Summary of study species and their climatic envelopes, listed from highest to lowest MAP for C_3_ and C4 plants. The climatic variables given are the mean, 5^th^ and 95^th^ percentiles of mean annual precipitation (MAP), and the mean, 5^th^ and 95^th^ percentiles of aridity index (AI), across the species’ distribution in Australia. Note that AI (Aridity Index) here is defined as mean annual potential evapotranspiration (PET) / mean annual precipitation (MAP); hence, a higher AI indicates a more arid environment.

|  | Binomial name | Tribe | Acronym | MAP (mm) | MAP<sub>Q5</sub> | MAP<sub>Q95</sub> | PET (mm) | AI | AI<sub>Q5</sub> | AI<sub>Q95</sub> |
| --- | --- | --- | --- | --- | --- | --- | --- | --- | --- | --- |
| C<sub>3</sub> | *Poa labillardierei* | Poeae | PL | 870 | 493 | 1436 | 1305 | 1.50 | 0.73 | 2.46 |
|  | *Microlaena stipoides* | Ehrharteae | MS | 848 | 555 | 1273 | 1306 | 1.54 | 0.89 | 2.31 |
|  | *Poa sieberiana* | Poeae | PS | 847 | 569 | 1250 | 1271 | 1.50 | 0.86 | 2.16 |
|  | *Austrostipa bigeniculata* | Stipeae | AB | 617 | 405 | 777 | 1283 | 2.08 | 1.49 | 3.40 |
|  | *Austrostipa scabra* | Stipeae | AS | 523 | 230 | 834 | 1731 | 3.31 | 1.59 | 6.99 |
|  | *Rytidosperma caespitosum* | Danthonieae | RC | 469 | 230 | 831 | 1656 | 3.53 | 1.40 | 6.74 |
|  | *Austrostipa elegantissima* | Stipeae | AE | 348 | 223 | 555 | 1615 | 4.64 | 2.33 | 7.32 |
| C<sub>4</sub> | *Themeda triandra* | Andropogoneae | TT | 799 | 429 | 1278 | 1582 | 1.98 | 1.02 | 3.38 |
|  | *Cymbopogon refractus* | Andropogoneae | CR | 777 | 508 | 1140 | 1515 | 1.95 | 1.2 | 2.81 |
|  | *Bothriochloa macra* | Andropogoneae | BM | 698 | 467 | 976 | 1424 | 2.04 | 1.33 | 3.06 |
|  | *Cynodon dactylon* | Cynodonteae | CD | 654 | 273 | 1176 | 1740 | 2.66 | 1.11 | 5.57 |
|  | *Dichanthium sericeum* | Andropogoneae | DS | 653 | 147 | 1116 | 2462 | 3.77 | 1.43 | 12.40 |
|  | *Chloris truncata* | Cynodonteae | CT | 543 | 270 | 839 | 1667 | 3.07 | 1.59 | 5.67 |
|  | *Enteropogon acicularis* | Cynodonteae | EA | 422 | 171 | 740 | 1929 | 4.57 | 2.05 | 10.85 |
|  | *Astrebla lappacea* | Cynodonteae | AL | 398 | 86 | 751 | 2762 | 6.94 | 2.18 | 20.59 |
|  | *Cenchrus ciliaris* | Paniceae | CC | 230 | 97 | 573 | 2275 | 9.89 | 3.21 | 18.53 |

### Study species

Sixteen common Australian grass species (seven C_3_ and nine C_4_ species) were used in this study (Table 1). Precipitation and temperature envelopes for the continental distribution of the species were obtained from the Worldclim database (https://www.worldclim.org/Fick & Hijmans, 2017) based on the distribution of the species in the Atlas of Living Australia (www.ala.org.au) using the “species map” package in R (https://remkoduursma.github.io/speciesmap). The distributions of the study species are shown in Fig. S1. The species all have sizeable ranges, but vary considerably in mean climate of origin, with species-average mean annual precipitation (MAP) ranging from 230 mm to 870 mm (Table 1). We also calculated a species mean aridity index (AI), using the inverse of the common definition, which we now calculate as the ratio of mean annual potential evapotranspiration (PET) to MAP. The re-definition of AI is done here so that high numbers indicate high aridity. This definition also facilitates our statistical analysis, because it gives a more even distribution of values across our species: values range from relatively low (1.50) to very high (9.89) aridity. Species were selected from a variety of tribes (four C_3_ tribes, three C_4_ tribes) to include diverse evolutionary backgrounds. All are perennial grasses that typically occur in unshaded habitats.

### Experimental design

Seeds were obtained from commercial seed suppliers (Nindethana seeds, Royston Petrie Seeds, and Austrahort Pty Ltd). After soaking in water overnight, seeds were sown on seed trays filled with commercial germination mix (Osmocote Seed raising and Cutting mix, Baulkham Hills, NSW 2153, Australia). After germination, seedlings were transferred to 30-litre square-faced plastic pots (Dimension: L x W x H= 290 mm x 290 mm x 390 mm, with holes at the bottom) filled with a sandy loam soil mixture. The soil mixture was obtained by mixing local (Yarramundi Paddock) soil (sieved to exclude plant debris) and commercial river sand in the ratio of 3:1 by mass. Eleven replicate pots were used for each of the study species, of which four were assigned to control (well-watered) and seven were assigned to the drought treatment (water-stressed). Six to eight individual grass plants were established in each pot, with the number of individuals being constant for each species.

Pots were placed inside a polytunnel. A randomized block design was adopted to account for environmental variation inside the polytunnel. Due to the time constraints associated with measurements, and the different seasonal requirements of the C_3_ and C_4_ species, the experiment was conducted in two sub-experiments matching the species groups to their main growing season: the first sub-experiment with nine warm-season C_4_ grass species ran from November 2019 to May 2020 (summer through autumn), while the second sub-experiment with seven cool-season C_3_ grass species ran from March 2020 to September 2020 (autumn through spring). Pots were spatially arranged into five blocks in sub-experiment 1 and four blocks in sub-experiment 2 with each block consisting of 20 pots representing different species and treatments. All replicates were maintained under well-watered conditions for three months. prior to pre-drought hardening. This involved irrigation being withheld for one week, then plants were irrigated back to field capacity and maintained at field capacity for two weeks (Duan et al., 2013). Using this technique, the plants experienced a moderate drought that initiated the physiological and biochemical ‘drought responses’ that conditioned the plants for the future drought, as would occur in a natural environment. An ongoing drought was then imposed by withholding irrigation. Drought periods lasted for 14 weeks (C_4_ grasses; sub-experiment 1) and 11 weeks (C_3_ grasses; sub-experiment 2).

Time-lapse cameras (phenocams) (WINGSCAPE/Moultrie) were placed above each block to capture the progression of leaf browning throughout the experiment. Photographs were taken four times each day, at 0800 h, 1000 h, 1400 h and 1600 h. One out of the four phenocam images was manually selected for green chromatic coordinate (GCC) calculation each day based on the contrast of the images. Each of the pots was assigned a separate region of interest. The GCC was calculated for each pot for each day using the *phenora* R package (https://bitbucket.org/remkoduursma/phenora/). To obtain relative GCC values (R_GCC_), GCC values were normalized to the observed minimum and maximum GCC values for each pot during the dry-down. The leaf water potentials corresponding to 12% (P_Br12_) and 50% (P_Br50_) reduction of leaf greenness were obtained by fitting a loess model function to values of R_GCC_ and leaf water potential using the *fitplc* R package (Duursma & Choat, 2017).

The hourly average volumetric soil water content (VSWC) of droughted pots was measured with 30 cm long soil moisture probes (CS650 TDR- connected to CR1000 data logger, Campbell Scientific Inc.) placed in the pots throughout the experiment. The VSWC of well-watered pots was measured using handheld TDRs on each gas exchange measurement date (CS650-30 cm). Environmental conditions (temperature, relative humidity and photosynthetic photon flux density (PPFD)) inside the polytunnel facility were recorded every 15 minutes by a data logger (CR300; Campbell Scientific Inc.) connected to a light sensor (Decagon Devices Inc., Pullman, WA, USA) and a temperature/relative humidity probe (HMP60-L, Campbell Scientific Inc., Logan, UT, USA) installed 1.5 m above the ground (Fig. S2).

### Gas exchange and leaf water potential measurements

Spot measurements of light-saturated photosynthesis (*A*_sat_), stomatal conductance (*g*_s_) and transpiration (*E*) were made throughout the dry-down phase of the experiment. Individual leaves used for gas exchange measurements were marked using a cotton thread knot. Measurements were made on 2-4 attached, recently fully-expanded leaves in the cuvette at a time, using identical portable open gas exchange systems (LI-6400XT, Li-Cor, Lincoln, NE, USA). Measurements were made with the fluorometer cuvette, which has a small chamber size (2 cm^2^) to maximize the proportion of cuvette occupied by leaves. Standard conditions maintained inside the cuvette were: reference [CO_2_] of 420 mmol mol^–1^, flow rate of 500 μmol s^-1^, block temperature of 25 ° C for C_4_ grasses and 20 ° C for C_3_ grasses, photosynthetic photon flux density (PPFD) of 1000 μmol m^–2^ s^–1^ and relative humidity in the range of 55 –65%. The PPFD level was determined to be saturating for all study species based on light response curves conducted before the dry-down was initiated (Fig. S3). Different standard temperatures were used for C_3_ and C_4_ species reflecting the different optimal temperature range for photosynthesis of these photosynthetic groups (Yamori et al., 2014). Parameters were logged when readings were visually stable, which was typically achieved within 4 to 6 minutes. All gas exchange parameters were expressed on a projected leaf area basis.

Leaves for predawn and midday leaf water potential measurements were collected between 0415 h and 0515 h and 1230 h and 1330 h, respectively. Leaves for predawn and midday leaf water potential measurements were taken from marked grasses. Leaf water potentials (Ψ_Predawn_ and Ψ_Midday_) were measured within 2-h after leaf collection using a Scholander-type pressure chamber with a maximum range of −10 MPa (PMS Instrument Company, OR, USA) on each gas exchange measurement date.

Maximum stomatal conductance (*g*_smax_) of study species was calculated as the mean of the maximum stomatal conductance of all eleven replicates of each species measured under well-watered conditions. Similarly, maximum photosynthetic rate (A_max_) of study species was also calculated. The stomatal closure traits (leaf water potentials inducing 12% (P_gs12_), 50% (P_gs50_) and 88% (P_gs88_) loss of stomatal conductance), were obtained using the “loess” model in the “fitcond” function in the fitplc package in R (Duursma & Choat, 2017) (Fig. S4). Similarly, leaf water potentials inducing 12% (P_A12_), 50% (P_A50_) and 88% (P_A88_) loss of photosynthetic rate were also obtained (Fig. S5).

Effective plant hydraulic conductance (*k*_p_, mmol m^-2^ s^-1^ MPa^-1^) for each of the gas exchange measurements was calculated using the following equation:

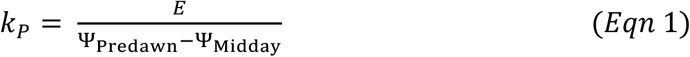

where *E* is transpiration rate per unit leaf area, Ψ_Predawn_ is predawn leaf water potential and Ψ_Midday_ is midday leaf water potential. Maximum effective plant hydraulic conductance (*k*_pmax_) of each study species was calculated as the mean of the maximum *k*_P_ value calculated for each replicate of that species. Similar to stomatal closure traits, hydraulic impairment traits for plant hydraulic conductance (P_kp12_ and P_kp50_) were also calculated (Fig. S6-7). Note that not all leaf water potential thresholds were able to be calculated for all study species. This principally affected four C_4_ species, namely *Astrebla lappacea, Cenchrus ciliaris, Cynodon dactylon* and *Enteropogon acicularis* (see Supplementary Figures for details).

### Pressure-volume curves

Pressure–volume curves were measured using the bench drying method (Schulte & Hinckley, 1985) for six replicate leaves per species. Leaves were sampled from well-watered individuals and rehydrated overnight. Images of rehydrated leaves were taken, and leaf area was calculated using ImageJ software (https://imagej.nih.gov/ij/). Leaves were cut near the leaf base before the first water potential measurement. In order to avoid leaf damage, leaves were wrapped in parafilm prior to measurements. Leaf and parafilm were weighed separately before wrapping. Immediately after wrapping, leaf water potential was measured using the pressure chamber. Following water potential measurements, leaf and parafilm were weighed. After each measurement, leaves were kept in a zipped aluminium foil bag to slow dehydration. This process was repeated until Ψ_leaf_ reached -3 to -3.5 MPa. Leaves were then oven-dried at 70°C for 48 hours and oven-dry weights were obtained. To obtain turgor loss point, pressure-volume curve analysis was performed using the R function developed by German Vargas (https://github.com/gevargu/). Capacitance values before and after turgor loss were calculated using the following equation:

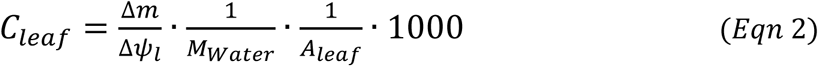

Where *C_leaf_* is the capacitance of the leaf (mmol MPa^-1^ m^-2^), Δ*m* is the change in the leaf fresh mass (g), Δ*ψ_l_*is the change in leaf water potential (MPa), *M_Water_* is the molecular mass of water (g mol^-1^) and *A_leaf_* is leaf area (m^2^).

### Leaf hydraulic conductance

The response of leaf hydraulic conductance to declining leaf water potential was measured using the rehydration kinetics method (RKM) (Brodribb & Holbrook, 2003). This method captures loss of both xylary and extra-xylary conductance and is commonly used for grasses (Corso et al., 2020; Griffin-Nolan et al., 2019; Holloway-Phillips & Brodribb, 2011; T. W. Ocheltree et al., 2016). Leaves used for hydraulic conductance measurements were obtained from well-watered individuals (either two leaves from the same tiller or adjacent leaves) and allowed to desiccate slowly for varying intervals until a wide range of leaf water potentials were achieved. Initial leaf water potential (Ψ_0_) was measured on the first leaf. Based on Ψ_0_ obtained for the first leaf, the second leaf was rehydrated for a pre-determined period (30-180 seconds). After rehydration, the second leaf was re-cut above the water line and placed in a pressure chamber to get final rehydrated leaf water potential (Ψ_f_). Leaf hydraulic conductance (*k*_l_) was calculated as follows:

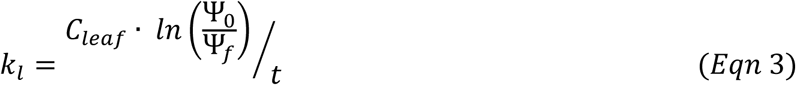

Where *C_leaf_* is the leaf capacitance (mmol MPa^-1^ m^-2^) calculated from pressure-volume analysis (using Equation 2), Ψ_0_ , Ψ*_f_* are initial and final rehydrated leaf water potentials and *t* is the rehydration time in seconds.

Maximum leaf hydraulic conductance (*k*_lmax_) was calculated as the mean of the six highest values of *k_l_* obtained for each species. Leaf hydraulic vulnerability curves were produced by plotting *k_l_* values against respective initial leaf water potential (Ψ_0_) values. As shown in Fig. S6, the leaf hydraulic impairment traits, defined as the leaf water potentials inducing 12% (P_kl12_), 50% (P_kl50_) and 88% (P_kl88_) loss of leaf hydraulic conductance, were obtained by fitting a loess model function to values of *k_l_*and Ψ_0_ values using the *fitplc* R package (Fig. S7) (Duursma & Choat, 2017).

### Data analysis

All statistical analyses were conducted using Rstudio (R v.4.0.3, RStudio 1.3.1093). Curve fittings of stomatal conductance, leaf hydraulic conductance, effective plant hydraulic conductance, photosynthesis and leaf greenness were performed using R with the *fitplc* package (Duursma & Choat, 2017). Example curves for each process, and the results of the fitting, are shown in Fig. 1. Linear regression (*lm* function) and Pearson correlation (*Corr* function) were used to assess correlations among stomatal closure, hydraulic impairment, and leaf browning, with photosynthetic and turgor loss traits of study species, and correlations of those traits with climatic variables.

**Fig. 1.**
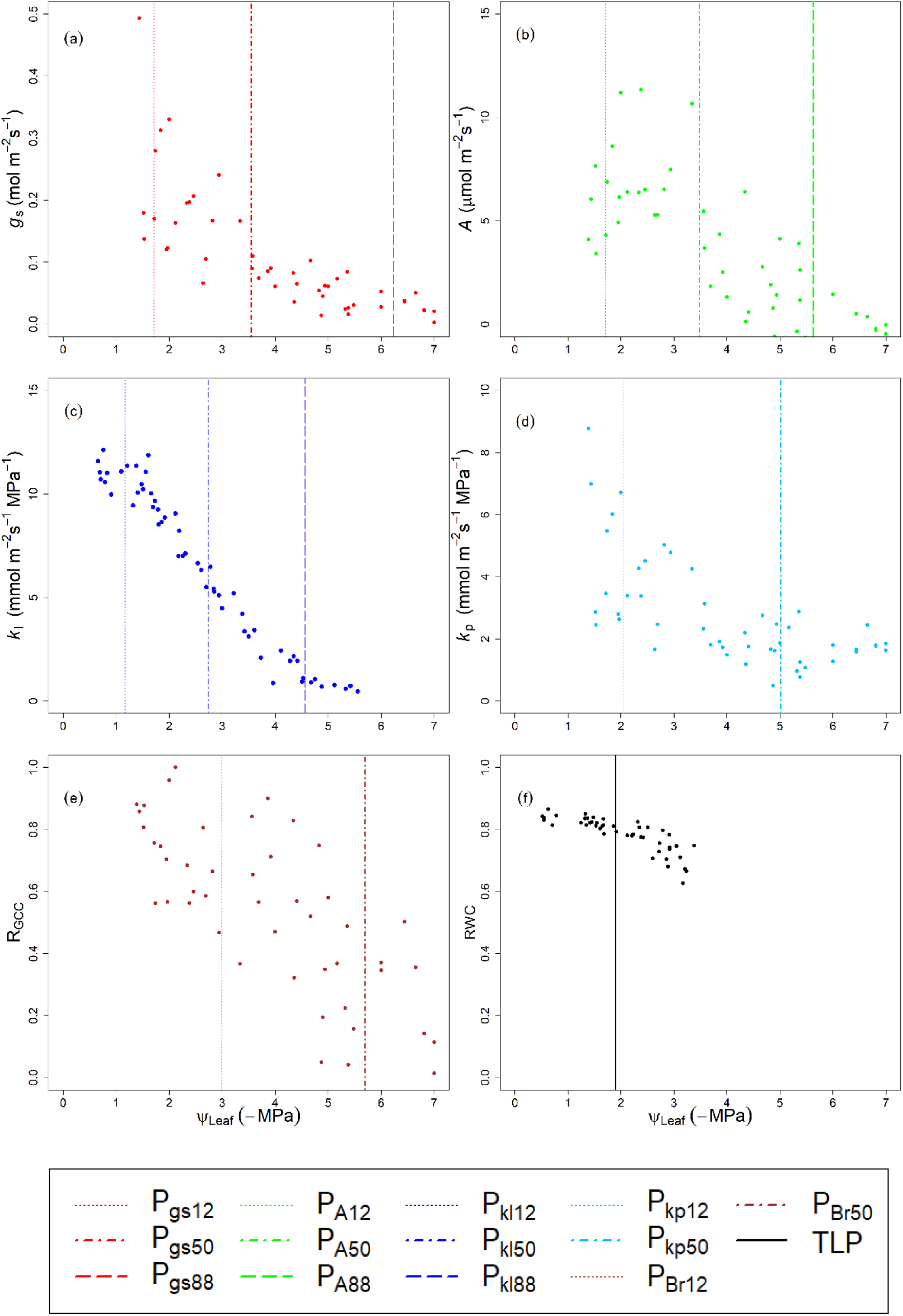
An example of data measured, for one species (*Poa labillardieri*) (a) Stomatal conductance (*g*_s_), vs midday leaf water potential, (b) Photosynthesis (*A*) vs midday leaf water potential, (c) Leaf hydraulic conductance (*k*_l_) vs initial leaf water potential (d) Effective plant hydraulic conductance (*k*_p_) vs midday leaf water potential, (e) Leaf greenness, relative to maximum and minimum (R_GCC_), vs midday leaf water potential, (f) Relative leaf water content (RWC) vs leaf water potential. Vertical lines in a-e indicate estimated % loss values for each metric: dotted lines indicate 12% loss, dot-dash 50% loss and dashed lines 88% loss. Vertical solid line in (f) indicates turgor loss point (TLP).

## Results

The leaf water potential thresholds for stomatal closure, loss of hydraulic conductance, turgor loss, impairment of photosynthesis and leaf browning are shown in Fig. 2. Within each of the C_3_ and C_4_ species groups, the variability across species in gas exchange, hydraulic impairment, leaf browning traits and turgor loss point is small. Most traits varied across species by less than 1.5 MPa. The largest ranges were for P_gs50_ and P_Br50_ in the C_4_ group, which varied by 2.18 and 2.33 MPa from most negative to the least negative, respectively.

**Fig. 2.**
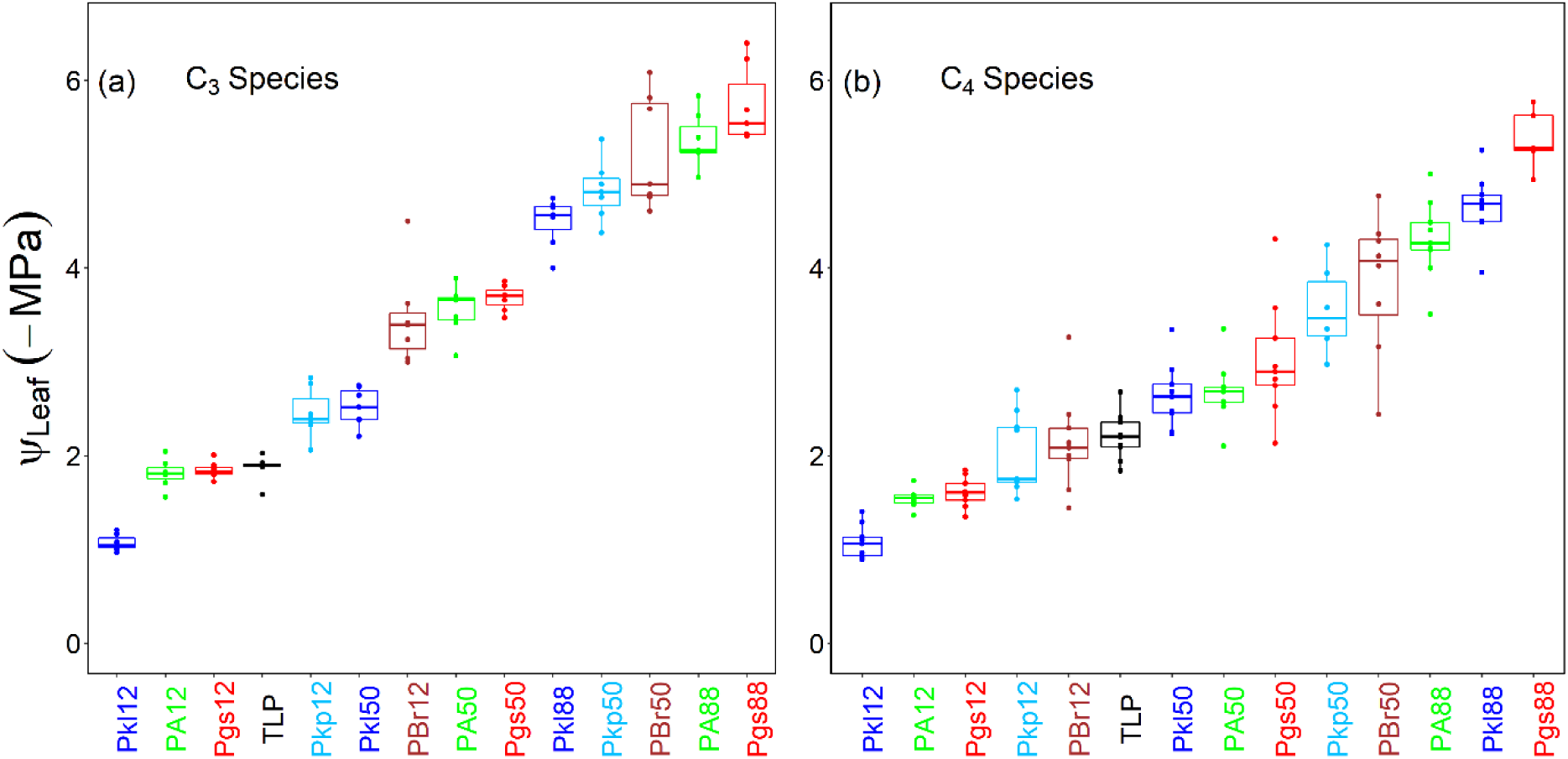
Sequence of stomatal closure, hydraulic impairment and leaf browning events in (a) C_3_ and (b) C_4_ species in response to declining leaf water potential during experimental dry-down. Dots indicate the individual trait values of either C_3_ or C_4_ species and grid boxes indicate 25^th^, 50^th^ (median) and 75^th^ percentile and inter-quartile range of the trait values within each photosynthetic group. Traits are ordered left to right for C_3_ and C_4_ species in the ascending order of the mean of each trait. Colours used for drought response traits relate to the graphs presented in Fig. 1. Number of observations (n) = 4-7 (C_3_), 7-9 (C_4_). To aid comparison between groups, confidence intervals for differences between C_3_ and C_4_ values are given in Table 2.

**Table 2.** Difference between C_3_ and C_4_ species for traits shown in Figure 2, including water potential thresholds for stomatal conductance (P_gs_), photosynthesis (P_A_), leaf hydraulic conductance (P_kl_), effective plant hydraulic conductance (P_kp_), leaf browning (P_Br_) and turgor loss point (TLP). Negative numbers indicate that C_3_ values are lower than values for C_4_ species.

| Metric | Percentage reduction | Mean difference | CI (2.5%, 97.5%) |
| --- | --- | --- | --- |
| P<sub>gs</sub> | 12 | **-0.18** | **(-0.36, 0.00)** |
|  | 50 | -0.36 | (-0.90, 0.18) |
|  | 88 | -0.21 | (-0.65, 0.22) |
| P<sub>A</sub> | 12 | **-0.26** | **(-0.40, -0.13)** |
|  | 50 | **-0.87** | **(-1.20, -0.54)** |
|  | 88 | **-1.05** | **(-1.45, -0.65)** |
| P<sub>kl</sub> | 12 | 0.00 | (-0.16, 0.16) |
|  | 50 | 0.12 | (-0.19, 0.43) |
|  | 88 | 0.16 | (-0.17, 0.50) |
| P<sub>kp</sub> | 12 | -0.16 | (-0.90, 0.62) |
|  | 50 | **-1.09** | **(-1.68, -0.51)** |
| P<sub>Br</sub> | 12 | **-0.66** | **(-1.13, -0.18)** |
|  | 50 | -1.48 | (-2.08, -0.89) |
| TLP |  | 0.33 | (0.10, 0.56) |

In both C_3_ and C_4_ species groups, the progression of stomatal closure, hydraulic impairment and leaf browning events occurred in tandem, with 12% loss of stomatal closure, hydraulic impairment, and reduction of canopy/leaf greenness, happening prior to the 50% loss events, which in turn occurred prior to the 88% loss events. Turgor loss occurred before P_gs50_ and P_kl50_ in both C_3_ and C_4_ functional groups, and P_gs88_ (stomatal closure) did not occur until after P_kl88_ and P_Br50_, i.e. after significant browning and loss of hydraulic conductance. Consequently, the HSM (P_gs88_ – P_kl50_) was negative for all C_3_ and C_4_ grass species, with a range of -2.65 to -3.75 MPa and -2.33 to -3.31 MPa, respectively. The values of P_kp50_ (loss of effective plant hydraulic conductance) were much lower than those of P_kl50_ (loss of leaf hydraulic conductance from rehydration kinetics method), by 1.65 MPa on average. However, P_gs88_ was lower than both, generating negative hydraulic safety margins (−3.15 MPa for P_kl50_, and -1.47 MPa for P_kp50_). This sequence contrasted strongly with the sequence of events that we hypothesized based on observations from trees.

There were relatively few differences in the sequence of events between C_3_ and C_4_ grass species (Fig. 2, Table 2). The chief difference was that the leaf browning events (P_Br12_ and P_Br50_) occurred at higher (less negative) leaf water potential in C_4_ species compared to C_3_ species (Table 2), such that P_Br12_ occurred after TLP in C_3_ species and before TLP in C_4_ species. In addition, in C_4_ grasses the decline in photosynthetic rate occurred at higher leaf water potentials (P_A12_, P_A50,_ P_A88_) and the decline was steeper (Fig. S5), relative to C_3_ grasses; however, the sequence in which these events occurred relative to the other processes did not differ between groups. There were also small differences in mean TLP, P_kp50_ and P_gs12_ between C_3_ and C_4_ species (Table 2).

In addition to differences in drought response traits, we also found a difference between C_3_ and C_4_ species in the relationship between predawn water potential (Ψ_Predawn_) and volumetric soil water content (VSWC) (Fig. 3). Compared to C_4_ grasses, the decline in predawn water potential of C_3_ species occurred at a higher VSWC, and the rate of decline was steeper. Thus, for a given VSWC, C_3_ species had lower (more negative) leaf water potentials than C_4_ species.

**Fig. 3.**
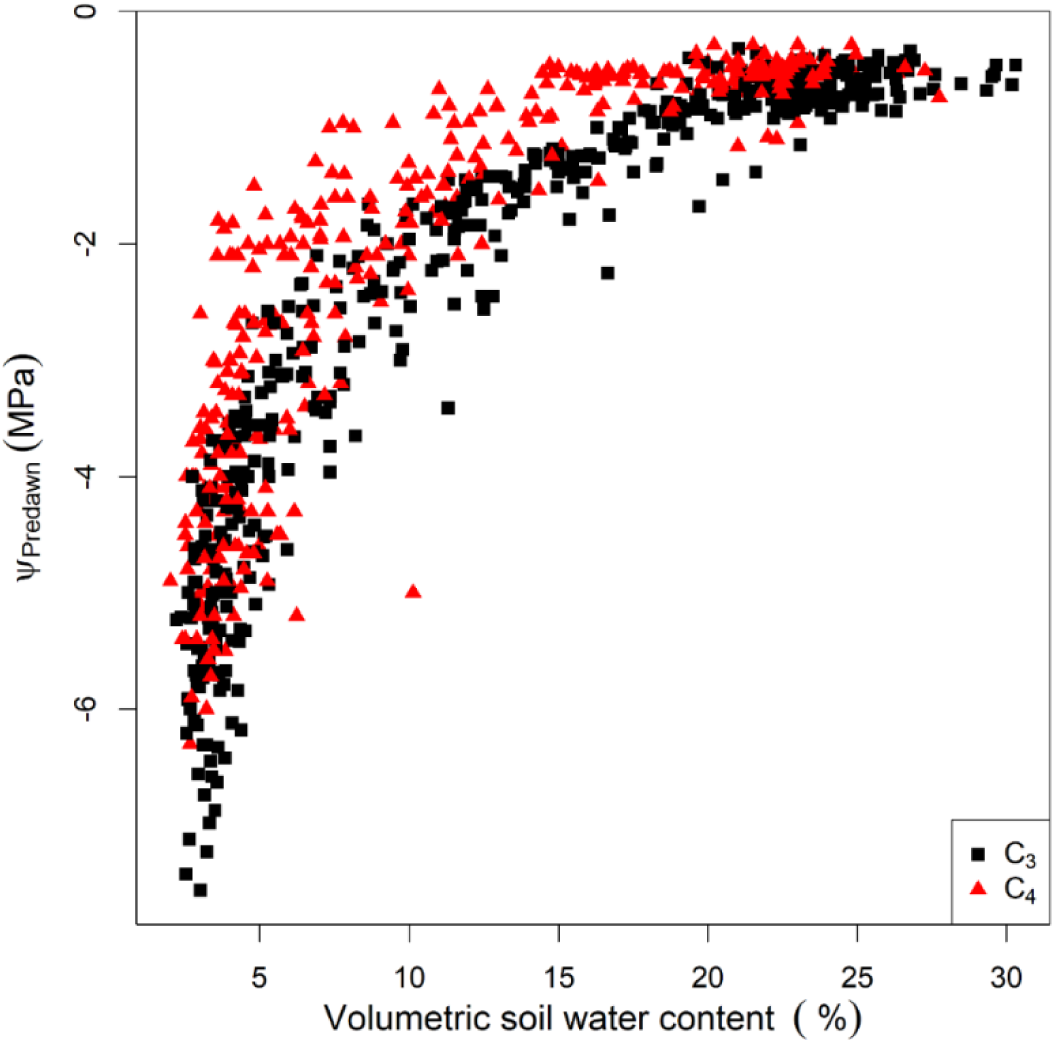
Decline in predawn leaf water potential versus volumetric soil water content in C_3_ and C_4_ grass species.

For trees, drought response traits are often correlated across species, indicating co-ordination between these traits. In Fig. 4, we examined correlations among these traits for the grass species. There was no significant correlation between P_kl50_ and P_gs50_, between P_kl50_ and P_Br50_, or between P_gs50_ and TLP (Fig. 4a, b and c). There was a significant correlation between P_gs50_ and P_A50_ (Fig. 4d), but this reflects the fact that both traits are derived from the same dataset (leaf gas exchange). Fig. 5 shows the pairwise correlations between traits using Pearson’s correlation coefficients (*r*) for C_3_ and C_4_ groups, separately. While there are some statistically significant correlations for each group, they are not consistent across groups. For example, there was a significant positive correlation between P_Br50_ and P_gs50_ with r = 0.68 for C_4_ species, but the same relationship for C_3_ species showed a non-significant negative correlation with r = -0.74. Hence, the data do not support co-ordination of these traits across species. Similarly, TLP was significantly correlated with P_Br50_, but this correlation was driven by the difference between C_3_ and C_4_ species groups, rather than consistent across-species relationships (Fig. S9).

**Fig. 4.**
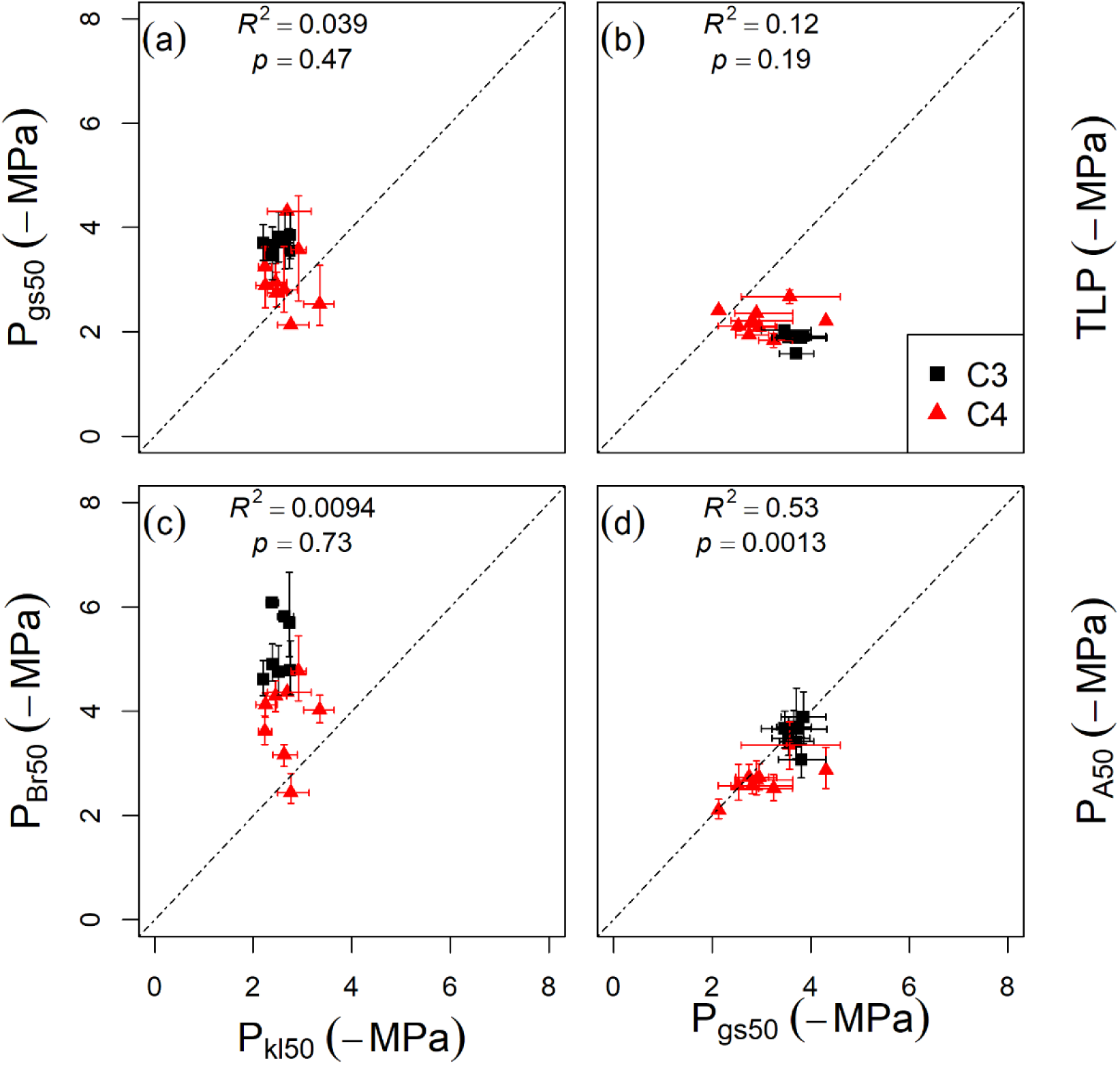
Relationships between (a) P_kl50_ and P_gs50_, (b) P_gs50_ and TLP (c) P_kl50_ and P_Br50_, and (d) P_kp50_ and P_Br50_ across all species, with 1:1 line drawn for comparison. Values of *R*^2^ and *p* for the relationships are given on each panel.

**Fig. 5.**
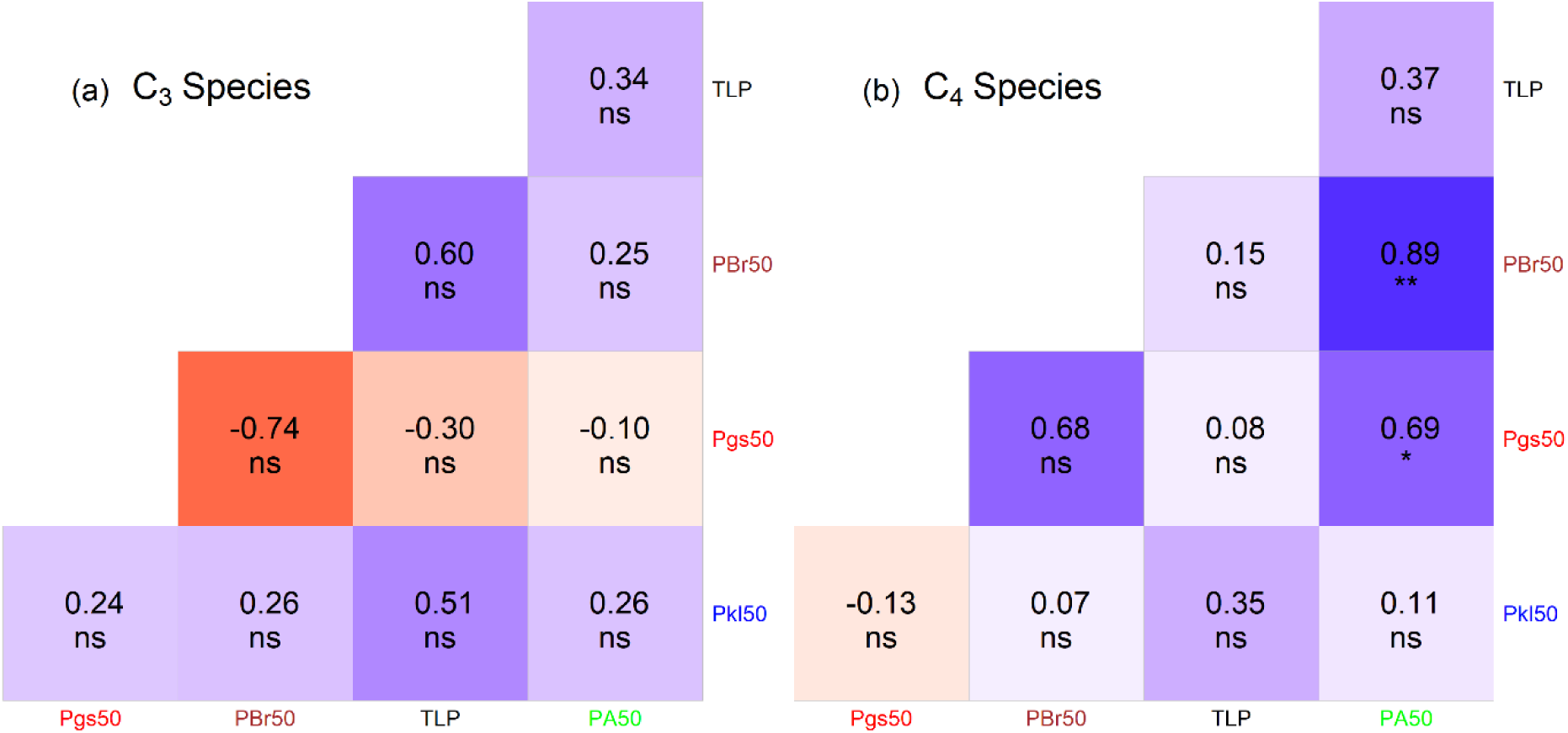
Pearson’s correlation coefficients (r) for relationships among stomatal closure, hydraulic impairment, and leaf browning traits for (a) C_3_ species and (b) C_4_ species. Statistical significance is indicated as follows: ns, non-significant; *, 0.01 < p < 0.05; **, p <0.01.

We tested correlations between stomatal closure, hydraulic impairment and leaf browning traits, and the climatic variables representing species’ climate of origin (Fig. 6). The climate variables examined include the mean MAP and AI across the species’ distributions, as well as the driest end of their distribution, represented by the driest percentiles (5^th^ and 95^th^) of annual precipitation (MAP_Q5_) and the aridity index (AI_Q95_), respectively. We found significant relationships between P_Br50_ and AI (Fig. 6k) and AI_Q95_ (Fig. 6l). However, these relationships did not meet expectations; the grass species originating from low rainfall environments (higher AI and AI_Q95_) showed higher P_Br50_ than the grass species originating from high rainfall environments. We did not find any significant relationships between other drought response traits (i.e. P_gs50_, P_kl50_, HSM and TLP) and the climate variables. Although there was a large range in the climatic variables (MAP: 230 – 870 mm yr^-1^, MAP_Q5_: 86 – 569 mm yr^-1^, AI: 1.50 – 9.89, AI_Q95_: 2.31 – 20.59), the variability in the drought response traits among study species was small, and appeared not to be related to climate.

**Fig. 6.**
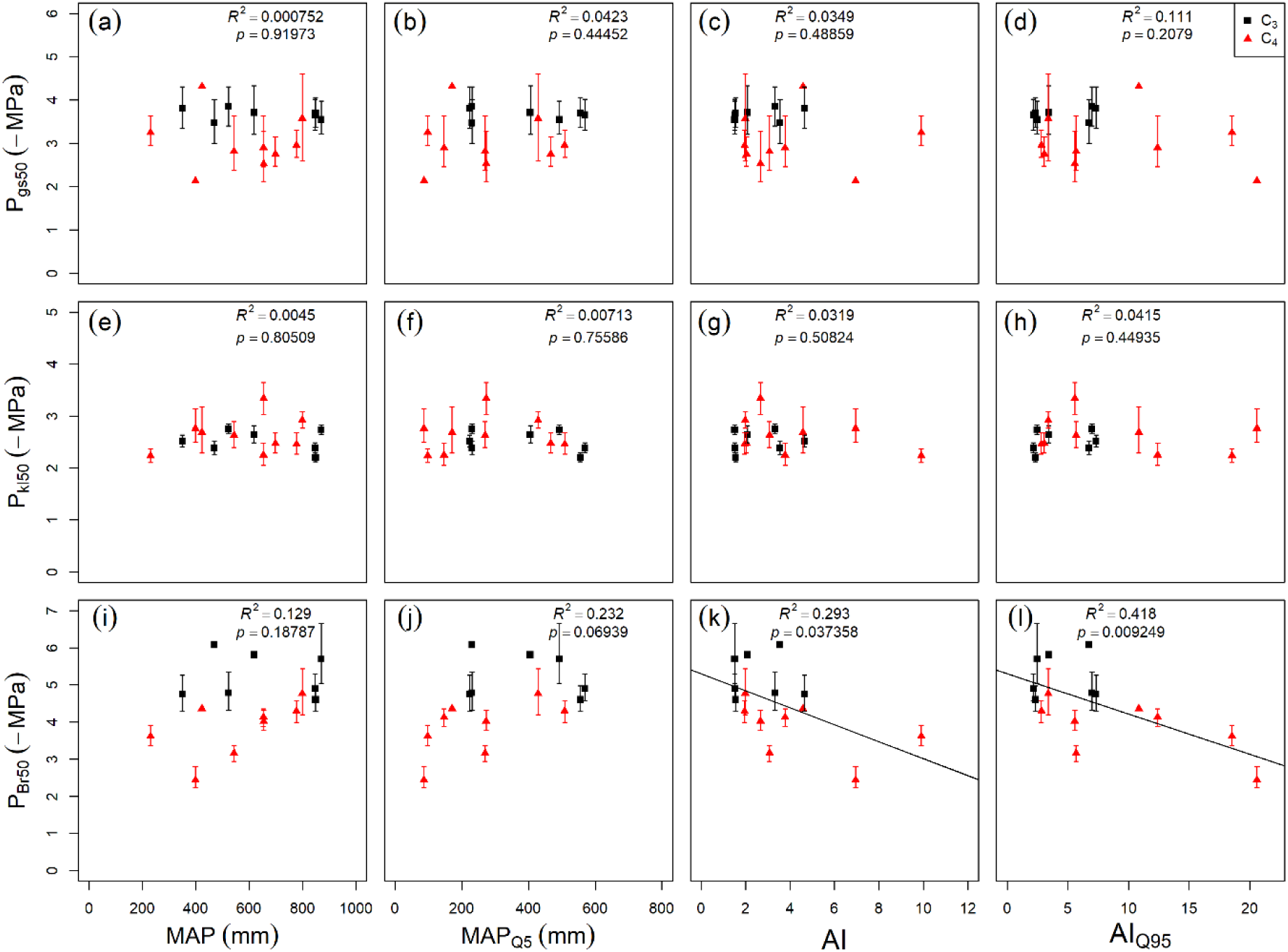
Trait correlations with climate of origin. Relationships between (a-d) stomatal closure (P_gs50_ - leaf water potential inducing 50% loss of stomatal conductance), (e-h) hydraulic impairment (P_kl50_ - leaf water potential inducing 50% loss of leaf hydraulic conductance), and (i-l) leaf browning (P_Br50_ - leaf water potential at 50 % reduction of leaf greenness) traits, with the climatic variables of mean annual precipitation (MAP), 5^th^ quantile of MAP (MAP_Q5_), aridity index (AI) and 95^th^ quantile of aridity index (AI_Q95_), where AI is defined as mean annual PET / MAP so that high values = high aridity. Regression lines indicate statistically significant relationships at p<0.05.

Finally, we tested for safety-efficiency trade-offs in maximum leaf hydraulic conductance (*k*_lmax_), maximum plant hydraulic conductance (*k*_pmax_) and maximum stomatal conductance (*g*_smax_) (Fig. 7). There was no evidence for a trade-off in *k*_lmax_ (Fig. 7a) or *k*_pmax_ (Fig. 7b), either within C_3_ or C_4_ groupings, or across both groups. There was a weak relationship between maximum stomatal conductance (*g*_smax_) and the leaf water potential at 50% loss of stomatal conductance (P_gs50_) (Fig. 7c). However, the relationship did not reflect a safety-efficiency trade-off and was more likely driven by differences between the C_3_ and C_4_ groupings, where C_3_ species tended to have both higher *g*_smax_ and more negative P_gs50_ than C_4_ species.

**Fig. 7.**
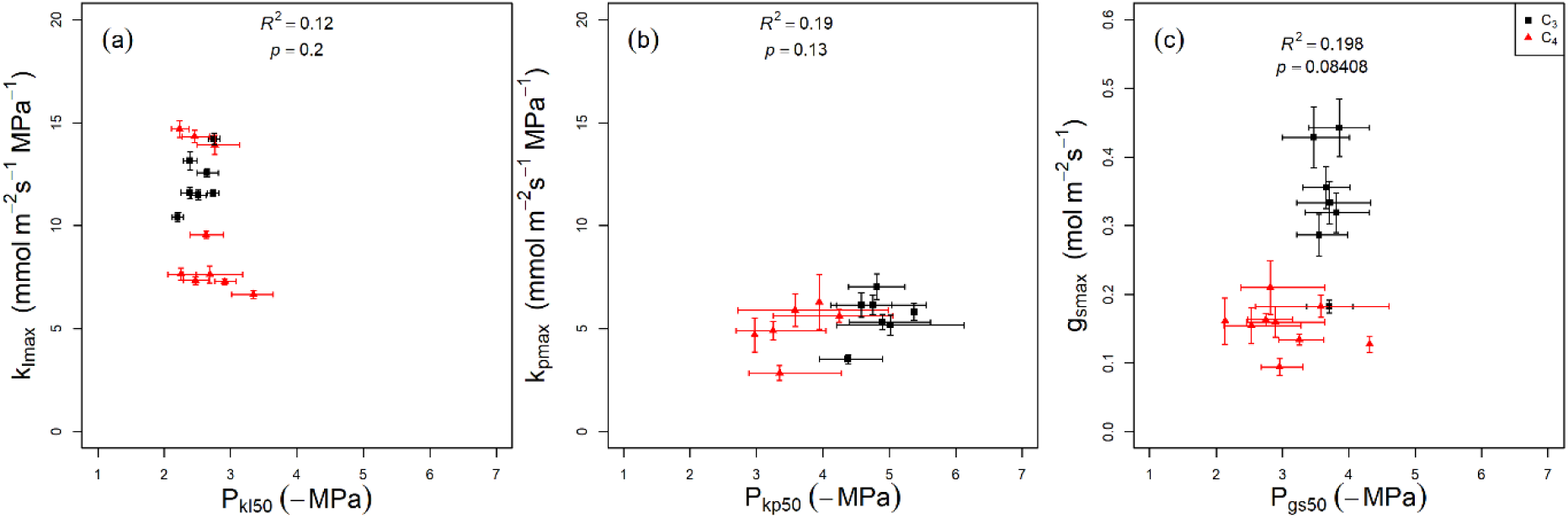
Safety-efficiency trade-offs for (a) leaf hydraulic conductance (K_l_), (b) plant hydraulic conductance (*K*_p_) and (c) stomatal conductance (*g*_s_). Each panel shows the relationship between the maximum rate of the process, and the water potential at 50% loss of that process.

## Discussion

We found that the sequence of drought response traits in grasses differed markedly from that typically observed in trees. Stomatal closure occurred concurrently with hydraulic impairment and leaf browning, in contrast to trees where stomatal closure typically occurs before the onset of hydraulic dysfunction and leaf loss (Blackman, Creek, et al., 2019; Brodribb & Jordan, 2008; Choat et al., 2018; Li et al., 2018). In addition, drought response traits showed no correlation with species’ climate of origin. The contrasting sequence and co-ordination of drought response traits in grasses relative to trees highlight the need for grass-specific frameworks to accurately represent drought responses in process-based models.

### Sequence of stomatal closure, hydraulic impairment and leaf browning events in grasses

A key finding was that stomata remained open well past the point of 50% loss of hydraulic conductance, implying that the grasses in our study had negative hydraulic safety margins. Our results are consistent with those of Holloway-Phillips & Brodribb (2011a), who reported simultaneous progression of both stomatal closure and leaf hydraulic impairment events in two C_3_ grasses (*Lolium multiflorum* and *Festuca arundinacea*), and those of Griffin-Nolan et al. (2023) who found P_gs50_ to occur well after P_kl50_ in 7 eastern Australian grasses. This finding for grasses contrasts strongly with trees under water-stressed conditions, in which stomatal closure (P_gs88_) typically precedes the impairment of stem hydraulic function (P_50 stem_) (Creek et al., 2018; Li et al., 2018; Skelton et al., 2021).

There are two complementary explanations for this contrast. The first relates to the replacement cost of xylem function in grasses vs. trees. The sequence of stomatal closure and hydraulic impairment events in trees can be understood as an adaptation to avoid the significant construction cost associated with re-establishing xylem function of tree stems after hydraulic disruption (Gauthey et al., 2022; Gleason et al., 2016). In contrast, the hydraulic architecture of grasses consists mainly of leaves, which have a much lower replacement cost than conductive vessels of tree stems, suggesting that drought-induced hydraulic impairment is less costly for grasses. This inference is also supported by the relatively low thresholds for the onset of leaf senescence in grasses. We found that leaf browning occurred concurrently with stomatal closure, particularly in C_4_ species. Woody species also have lower thresholds for conductivity loss in leaves than in stems (Bartlett et al., 2016), leading to the “hydraulic fuse” hypothesis, that trees shed leaves to avoid loss of xylem function. However, leaf drop in trees typically does not occur until well after stomatal closure and the onset of stem embolism (Blackman, Creek, et al., 2019; Wolf et al., 2016). In contrast, foliar senescence is a common drought escape strategy in grasses originating from seasonally dry grasslands (Balachowski et al., 2016; T. W. Ocheltree et al., 2020; VanderWeide et al., 2014). While trees appear to prioritize protection of the hydraulic system by closing stomata to reduce water use, in grasses it may be more beneficial to keep stomata open beyond the onset of hydraulic impairment and instead reduce water use through reductions in leaf area through selective leaf senescence.

Secondly, the role of extra-xylary hydraulic conductance is important in understanding the distinctive functionality of grasses under drought. Previous studies of the impairment of leaf hydraulic conductance (*k*_l_) in grasses and other angiosperms under water-deficit have shown that significant reduction in *k*_l_ during the early stage of water-deficit is mainly caused by a decline in extra-xylary hydraulic conductance rather than by cavitation in leaf veins (Corso et al., 2020; T. W. Ocheltree et al., 2020; Scoffoni et al., 2017, 2023). The method used here to measure *k*_l_, the rehydration kinetics method (RKM), captures both extra-xylary and cavitation-induced declines in hydraulic conductance (Scoffoni et al., 2017). Corso et al. (2020) compared RKM with an optical method to measure xylem cavitation and showed that a decline in extra-xylary conductance contributed to the significant loss of *k*_l_ prior to cavitation in wheat. Similarly, our results can be compared with those of Jacob et al. (2022) who examined the hydraulic vulnerability of five pasture grass species in Australia, including *Themeda triandra*, using the optical hydraulic vulnerability technique which monitors xylem embolism. They found P_kl50_ values of C_3_ and C_4_ grasses ranged from -4.4 MPa to -6.08 MPa, considerably more negative than values observed here, with P_kl50_ of *Themeda triandra* as low as -6.08 MPa. In contrast, values for *Themeda triandra* using the RKM method include -2.91 MPa in the current study, and -1.45 MPa in the study by Griffin-Nolan et al. (2023). This comparison suggests a strong role for the extra-xylary hydraulic pathway in leaf hydraulic conductance of grasses, consistent with previous studies (Corso et al., 2020; T. W. Ocheltree et al., 2020; Scoffoni et al., 2017). Impairment of this pathway is likely to be considerably less costly to a plant than loss of xylary conductance, helping to explain why stomata remain open past the onset of conductance loss.

### Comparison between C_3_ and C_4_ species

Drought responses in grass species need to consider the effect of photosynthetic pathway, owing to the important differences in water relations (Ghannoum, 2008). Here we found, as expected, lower maximum stomatal conductance in C_4_ than C_3_ grasses (Fig. 7). However, we did not find a significant effect of photosynthetic pathway on stomatal or hydraulic drought response traits (Fig. 2, Table 2). Instead, the major differences we observed included (i) higher predawn leaf water potential for a given volumetric soil water content (Fig. 3); and (ii) stronger sensitivity of photosynthesis and leaf browning to leaf water potential in C_4_ species, relative to C_3_ species (Fig. 2, Table 2).

The higher predawn water potential in C_4_ grasses was unexpected, given the widespread assumption that Ψ_Predawn_ reflects soil moisture content (Drake et al., 2017). We considered whether this could be an artefact of conducting the C_3_ and C_4_ grass experiments at different times of the year (winter-spring for C_3_, summer-autumn for C_4_) (Fig. S2). However, this difference in timing might be expected to yield a lower Ψ_Predawn_ at a given VSWC in C_4_ species than C_3_ species, owing to the higher air temperature and VPD during the C_4_ experiment. Differences in Ψ_Predawn_ among species are also often attributed to differences in rooting depth (Bucci et al., 2009). However, such differences are unlikely to emerge in plants growing in 30-L pots. Instead, the difference could indicate differing capacities for overnight refilling of plant capacitance, with C_4_ grasses having more complete refilling than C_3_ grasses (Harrison Day et al., 2024; Stewart et al., 2025). This explanation would be consistent with work by O’Keefe & Nippert (2018), who showed higher rates of nocturnal transpiration in C_4_ species than C_3_ species. Our results are also consistent with those of Taylor et al. (2011), who observed more negative operating leaf water potentials for C_3_ than C_4_ grasses at similar soil moisture content.

Our finding that the drought response of photosynthesis occurs at higher leaf water potentials C_4_ grasses had a less negative P_A50_ than C_3_ grasses, indicating greater drought sensitivity of photosynthesis in C4 species, consistent with Ripley et al. (2010). However, comparison with previous studies requires some care in interpretation. Taylor et al. (2011) found greater stomatal drought sensitivity in C_3_ than C_4_ plants, but the analysis of Taylor et al. (2011) was based on soil moisture content rather than plant water potential, making direct comparisons difficult. In a large trait-based study of 426 grass species, Craine et al. (2013) found that Ψ_crit_, defined as the leaf water potential at the point of stomatal closure, did not differ on average between C_3_ and C_4_ groups, which is consistent with our finding for P_gs88_. We also observed a difference in the onset of browning between C_3_ and C_4_ grasses, with C_4_ grasses having an earlier onset of browning as measured by leaf water potential (Fig. S8). This finding accords with the slightly higher drought-induced leaf mortality of C_4_, compared to C_3_, grasses reported by Taylor et al. (2011).

While photosynthetic pathway appears to be a key determinant of some traits, several authors have highlighted that there are also large differences in functional traits among lineages within the grasses. For example, Donnelly et al. (2023) found lineage to be a major predictor for functional traits such as maximum plant height and foliar C:N. Our experiment was not explicitly designed to test for differences among lineages and we had only 1-4 species per lineage. Nonetheless, visualization of the dataset by lineage (Fig. S10) does not show any taxonomic clustering of trait values, indicating that lineage is likely not a better predictor than photosynthetic pathway for the traits considered here.

### Adaptation to climate of origin

Another key finding of this study was the lack of co-ordination among the drought response traits among species, and between the drought response traits and the climate of origin. These findings contrast strongly with the correlation commonly reported between stomatal closure and hydraulic impairment traits of tree species and their climate of origin (Blackman et al., 2014; Gleason et al., 2016; Li et al., 2018; Nardini & Luglio, 2014; Peters et al., 2021). It is, however, consistent with other literature finding relatively weak relationships between grass traits and climate of origin (Baird et al., 2025; Craine et al., 2013). Notably, the grass species used in this study overlap strongly geographically with the tree species studied by Li et al. (2018), and cover a similar range in MAP across species (188-1,125 mm in Li et al. (2018) and 230-870 mm in this study), yet the strong trait-climate relationships reported for trees in Li et al. (2018) were absent in our grass species. This highlights a clear difference in adaptive strategy between the two growth forms.

Notably, we found a relatively small range in measured P_kl50_ (−2.20 MPa to -3.34 MPa) across species, suggesting low adaptive variation in this trait (Fig. 4a). Other studies have found a wider range: for example, Ocheltree *et al*. (2016) reported a wider range of -1.80 MPa to - 4.78 MPa for P_kl50_ for nine C_4_ grasses and subsequently observed coordination between P_kl50_ and P_gs50_. Their method for P_kl50_ differed somewhat from ours: they measured *K*_leaf_ on leaves collected at different timepoints during a dry-down, while we measured leaf hydraulic conductance at declining leaf water potentials for bench-dried leaves, which were collected from well-watered grass individuals. It is possible that the different methodology leads to a stronger variation in P_kl50_ and a stronger relationship with P_gs50_. Nonetheless, similar to our finding, their results do not show clear adaptive variation, with P_kl50_ found to be poorly correlated with climate of origin.

One caveat of our study is that the grass species we studied have quite large geographic ranges (Fig. S1), and the climate metrics characterizing their distributions in Australia may not reflect the particular provenances used in this study. Traits may vary within species depending on provenance: for example, Jacob et al. (2025) shows variation in P_gs90_ and TLP with summer precipitation of origin among provenances of *T. triandra*. Unfortunately, the available provenance information for the seeds used in this study was too coarse to test for relationships with provenance climate of origin. However, Griffin-Nolan et al. (2023) examined a similar suite of species with plant material collected in the field at four locations ranging in AI from 3.2 to 8.9, including different provenances of the same species, and found no relationships with climate of origin, similar to our study.

The lack of relationship between drought response traits and climate of origin suggests that rather than adapting stomatal function and hydraulic conductance, grass species adapt to dry environments through alternative mechanisms. For example, dry-adapted grass species may have larger allocation of photosynthates to below-ground tissues, such as deep-rooted systems that enable them to acquire more water, or root crowns that stored nutrients for subsequent recovery (Brunner et al., 2015; Chandregowda et al., 2022; Kleczewski et al., 2012). Another major mechanism for adaptation to dry conditions in grasses is to shift phenological timing to match photosynthetic activity with water availability (Hufkens et al., 2016).

### Hydraulic safety and efficiency trade-off in grass leaves

We hypothesized a safety vs. efficiency trade-off in the hydraulic pathway of grasses, based on previous studies that found this trade-off in grasses (T. W. Ocheltree et al., 2016) and other plant functional types (Johnson et al., 2012; Nardini et al., 2012). However, we found no evidence for this trade-off in grass leaves. The theory behind this trade-off relates to the architecture of the xylem transport pathway: it is argued that the large pores in bordered pits required for high water transport rates are also more prone to cavitation. It is possible that our inability to separate the vascular (xylary) and extra-vascular pathways of the leaves (Nardini et al., 2010; T. W. Ocheltree et al., 2016; Sack et al., 2003; Scoffoni et al., 2017) may have limited our capacity to test for hydraulic trade-offs, since similar constraints may not apply to the extra-vascular pathway. However, our results are consistent with previous studies which have reported a lack of hydraulic safety and efficiency trade-off in leaves of trees (Blackman et al., 2010, 2014; Nardini & Luglio, 2014) and grasses (Jacob et al. 2022).

### Conclusion: Implications for modelling drought responses of grasses

Our ultimate goal with this work was to inform the mechanistic, process-oriented models needed to predict future drought vulnerability in terrestrial ecosystems. A number of such models incorporating stomatal and hydraulic traits have been developed for woody species (De Kauwe et al., 2020; Smith et al., 2014; Sperry et al., 2017; Tuzet et al., 2003). Many of these models use optimization approaches (Lu et al., 2016; Sperry et al., 2017; Venturas et al., 2018; Wolf et al., 2016), in which stomatal responses of plant species to drought are predicted from a trade-off between photosynthetic gain and hydraulic cost, with stomatal closure predicted to occur to avoid large hydraulic costs. However, our observations suggest that grasses do not optimize their stomatal behaviour in order to avoid costs associated with xylem embolism, as is the case with trees, and thus the optimization models are likely inappropriate to predict stomatal behaviour in grasses. Instead, drought responses are characterized by near-simultaneous declines in leaf gas exchange, leaf hydraulic conductance, including extra-xylary conductance, and green leaf area. The observation that thresholds for these processes are similar across species regardless of climate of origin is useful for modelling as it suggests that only a single set of parameter values for these processes is needed to characterize grassland drought responses. This study provides a useful empirical basis for deriving these parameterisations.

## Data Availability

The data used in this manuscript are publicly available at dx.doi.org/10.6084/m9.figshare.23538600

## Supplementary Information

The following **Supplementary** Information is available for this article:

**Fig. S1** Distribution of the grass species and the mean annual precipitation of their climate origin.

**Fig. S2** Variation in daily maximum, minimum and mean air temperature (T_air_), Vapour pressure deficit (VPD) and daily maximum photosynthetic photon flux density (PPFD) inside the Polytunnel facility during sub-experiment 1 and 2.

**Fig. S3** Light response curves of photosynthesis for studied C_3_ and C_4_ grass species

**Fig. S4** Loess fits for stomatal conductance (P_gs12,_ P_gs50_, P_gs88_).

**Fig. S5** Loess fits for photosynthetic rates of study species to respective midday leaf water potential during experimental dry-down.

**Fig. S6** Loess fits for effective plant hydraulic conductance (P_kp12_, P_kp50_, P_kp88_)

**Fig. S7** Loess fits for leaf hydraulic conductance (P_kl12_, P_kl50_, P_kl88_)

**Fig. S8** Loess fits for leaf browning (P_Br12_, P_Br50_)

**Fig. S9** Relationships between turgor loss point (TLP) and (a) leaf hydraulic impairment (P_kl50_) (b) plant hydraulic impairment (P_kp50_) (c) stomatal closure (P_gs50_) traits and (d) leaf browning (P_Br50_) traits.

**Fig. S10** Same as Figure 4, but with points colored to indicate grass tribe of each species.

**Table S1.** Table of symbols, with units and their definitions

