## Supplementary Tables and Figures for "Lack of co-ordination of stomatal, hydraulic and leaf browning traits in 16 perennial Australian grass species of differing climate origins"

### Supporting information

The following Supporting Information is available for this article:

**Fig. S1** Distribution of the grass species and the mean annual precipitation of their climate origin.

**Fig. S2** Variation in daily maximum, minimum and mean air temperature ($T_{air}$), Vapour pressure deficit (VPD) and daily maximum photosynthetic photon flux density (PPFD) inside the Polytunnel facility during sub-experiment 1 and 2.

**Fig. S3** Light response curves of photosynthesis for studied $C_3$ and $C_4$ grass species

**Fig. S4** Loess fits for stomatal conductance ($P_{gs12}$, $P_{gs50}$, $P_{gs88}$).

**Fig. S5** Loess fits for photosynthetic rates of study species to respective midday leaf water potential during experimental dry-down.

**Fig. S6** Loess fits for effective plant hydraulic conductance ($P_{kp12}$, $P_{kp50}$, $P_{kp88}$)

**Fig. S7** Loess fits for leaf hydraulic conductance ($P_{kl12}$, $P_{kl50}$, $P_{kl88}$)

**Fig. S8** Loess fits for leaf browning ($P_{Br12}$, $P_{Br50}$)

**Fig. S9** Relationships between turgor loss point (TLP) and (a) leaf hydraulic impairment ($P_{kl50}$) (b) plant hydraulic impairment ($P_{kp50}$) (c) stomatal closure ($P_{gs50}$) traits and (d) leaf browning ($P_{Br50}$) traits.

**Fig. S10** Same as Figure 4, but with points coloured to indicate grass tribe of each species.

**Table S1.** Table of symbols, with units and their definitions

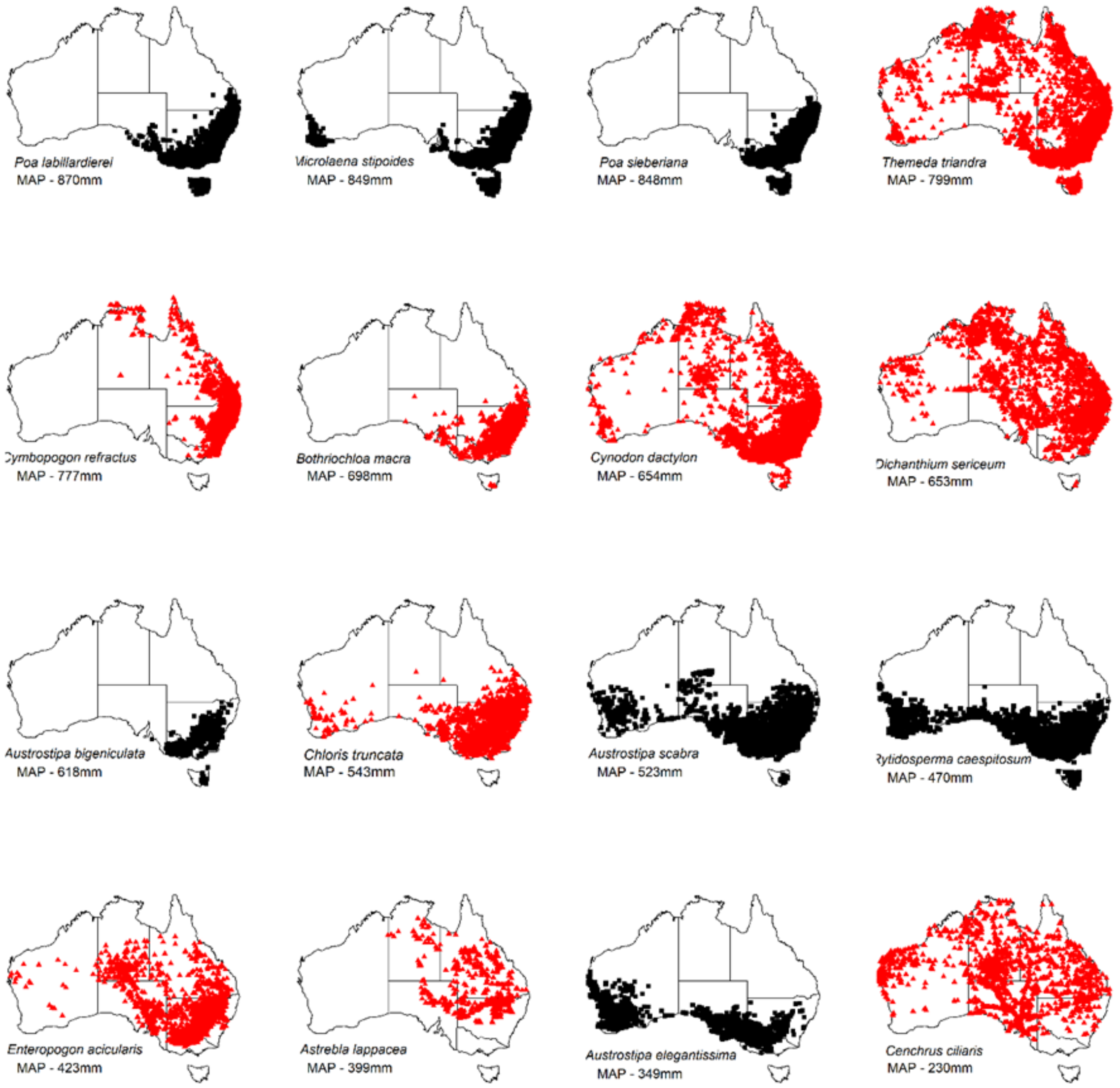


**Fig. S1** Distribution of the grass species and the mean annual precipitation of their climate origin. Species distribution information was obtained from the Atlas of Living Australia (https://www.ala.org.au/) using the "species map" R package developed by Remko Duursma. MAP values were calculated using the long-term rainfall data of each location obtained from Worldclim database. Black and red colours indicate $C_3$ and $C_4$ species, respectively.

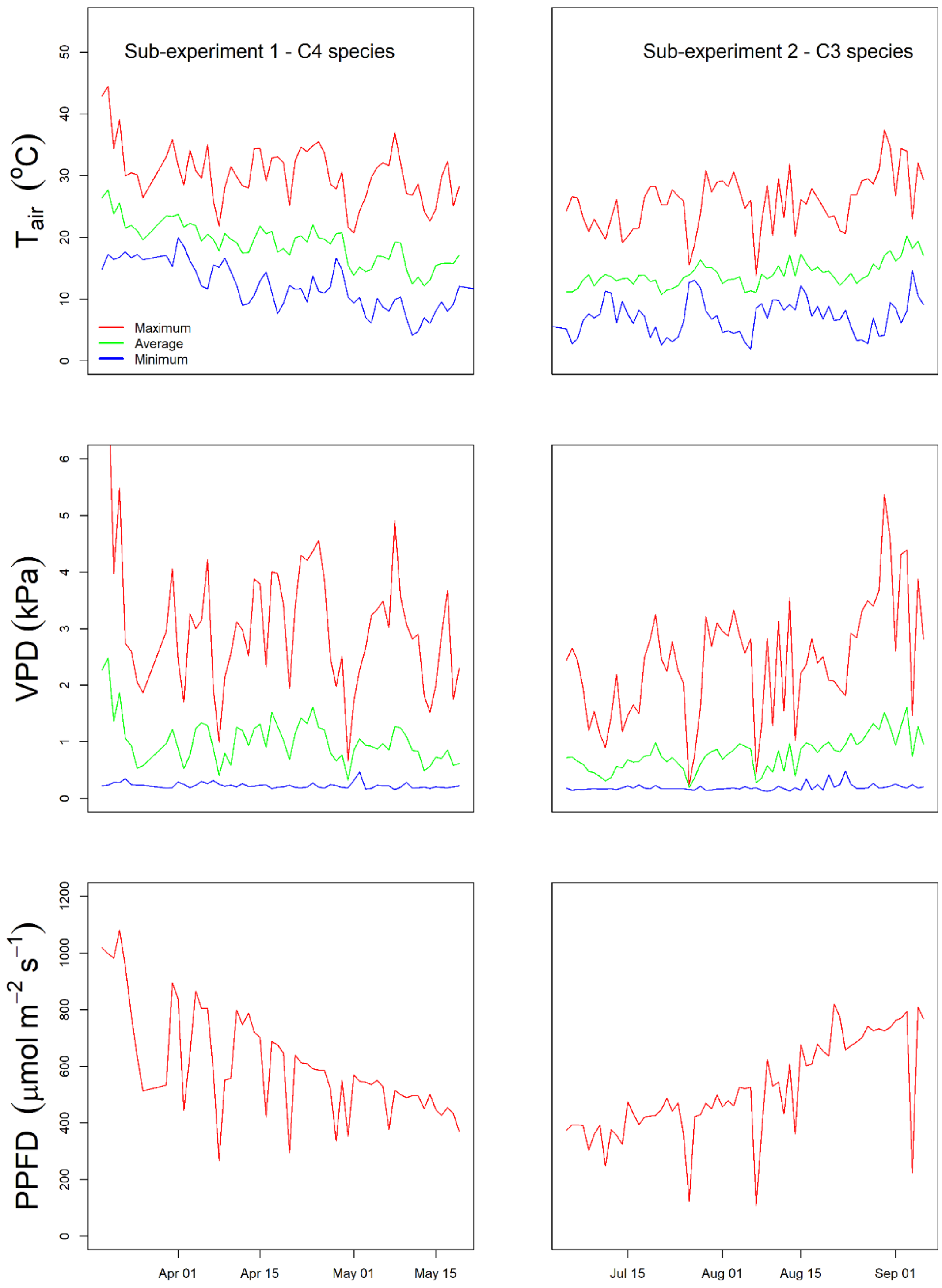


**Fig. S2** Variation in daily maximum, minimum and mean air temperature ($T_{air}$), Vapour pressure deficit (VPD) and daily maximum photosynthetic photon flux density (PPFD) inside the polytunnel facility during sub-experiment 1 and 2. Sub-experiment 1 with 9 $C_4$ grass species ran from November 2019 to May 2020 and Sub-experiment 2 with 7 $C_3$ grass species ran from March 2020 to September 2020

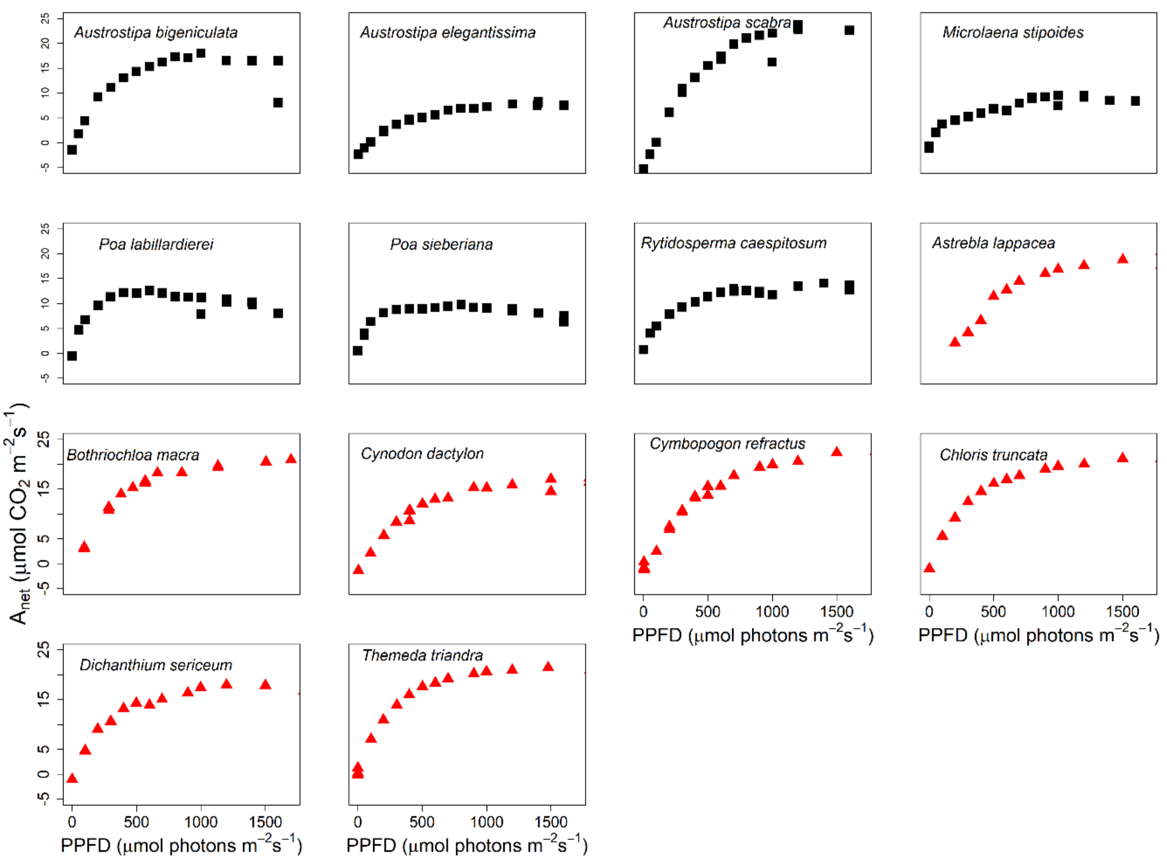


**Fig. S3** The light response of photosynthesis was measured in each study species at the outset of the experiment to determine an appropriate light level for leaf gas exchange measurements. For each species measurements were obtained from three well-watered replicates at light levels of (0, 50, 100, 200, 300, 400, 500, 600, 700, 800, 900, 1000, 1200, 1400 and 1600 µmol m–2 s–1) using LI-6400XT gas exchange measurement systems.

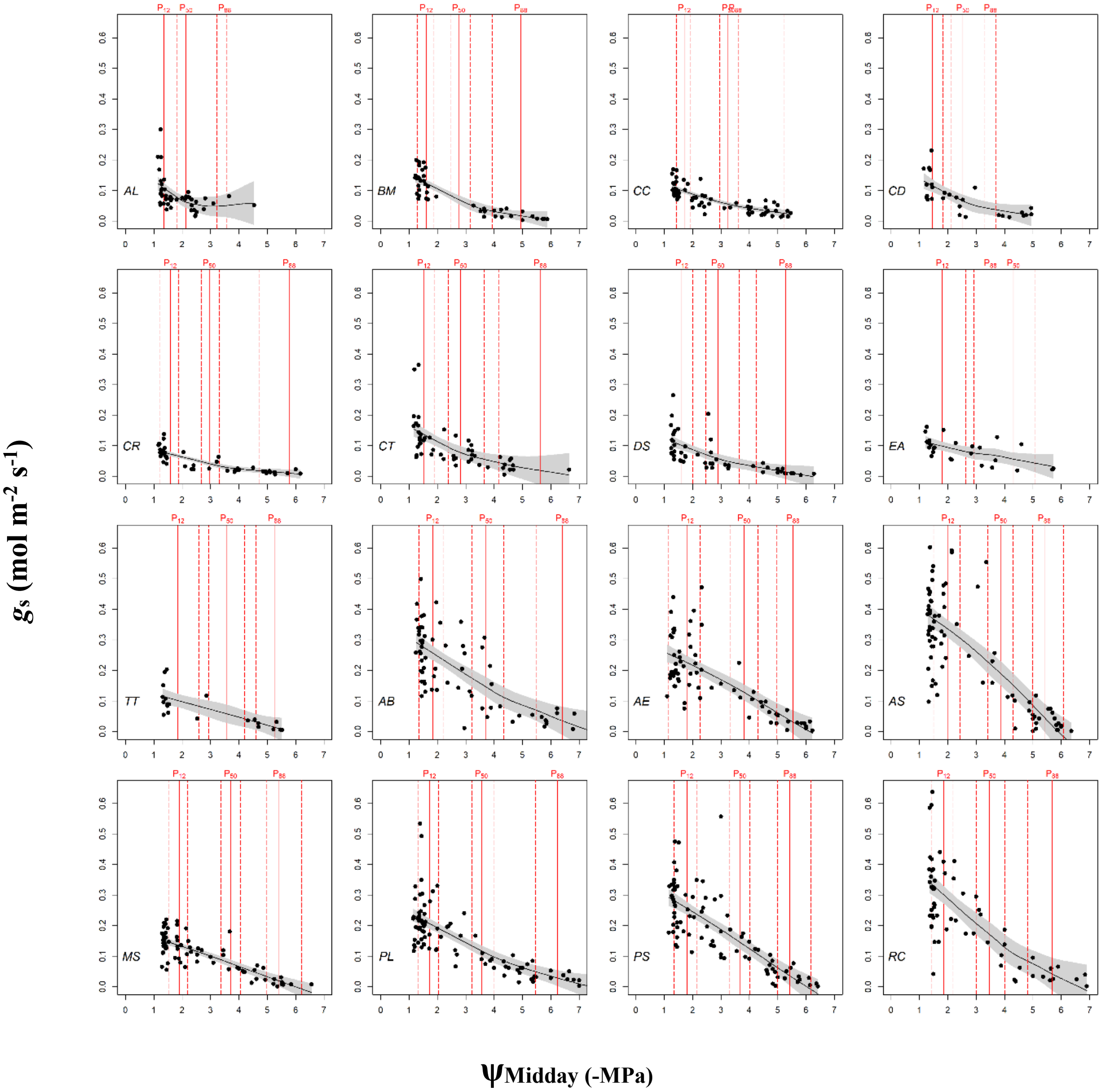


**Fig. S4** Loess fits for stomatal conductance ($P_{gs12}$, $P_{gs50}$, $P_{gs88}$) using fitcond function in fitplc package in R. Species abbreviations given in Table 1. For four species (AL, CC, CD and EA) Pgs88 values could not be fit. Red vertical lines show the fitted parameter value and confidence interval (CI) (2.5% and 97.5 %).

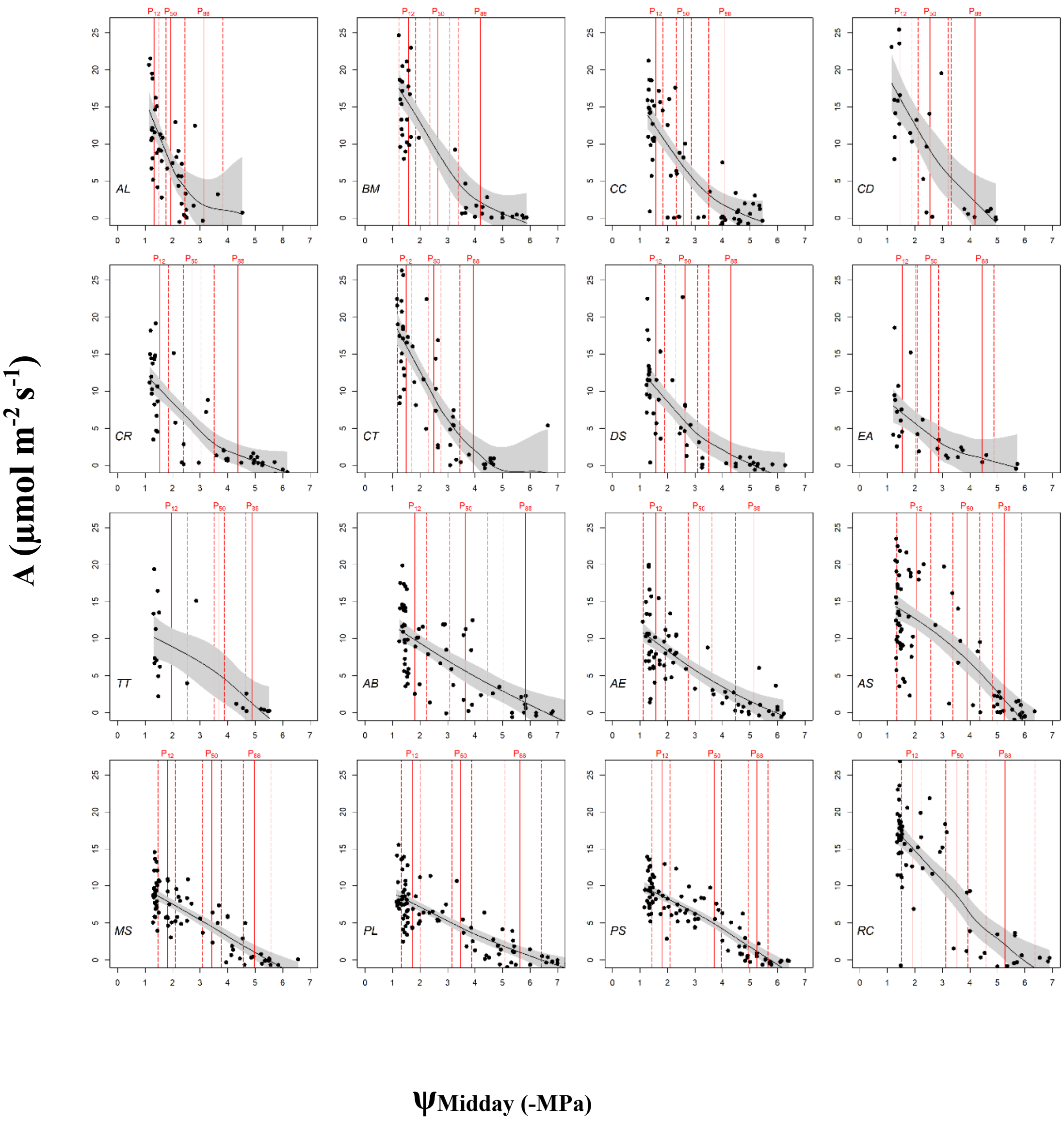


**Fig. S5** Loess fits for photosynthetic rates of grass species to respective midday leaf water potential during experimental dry-down. Red vertical lines show the fitted parameter value and confidence interval (CI) (2.5% and 97.5 %).

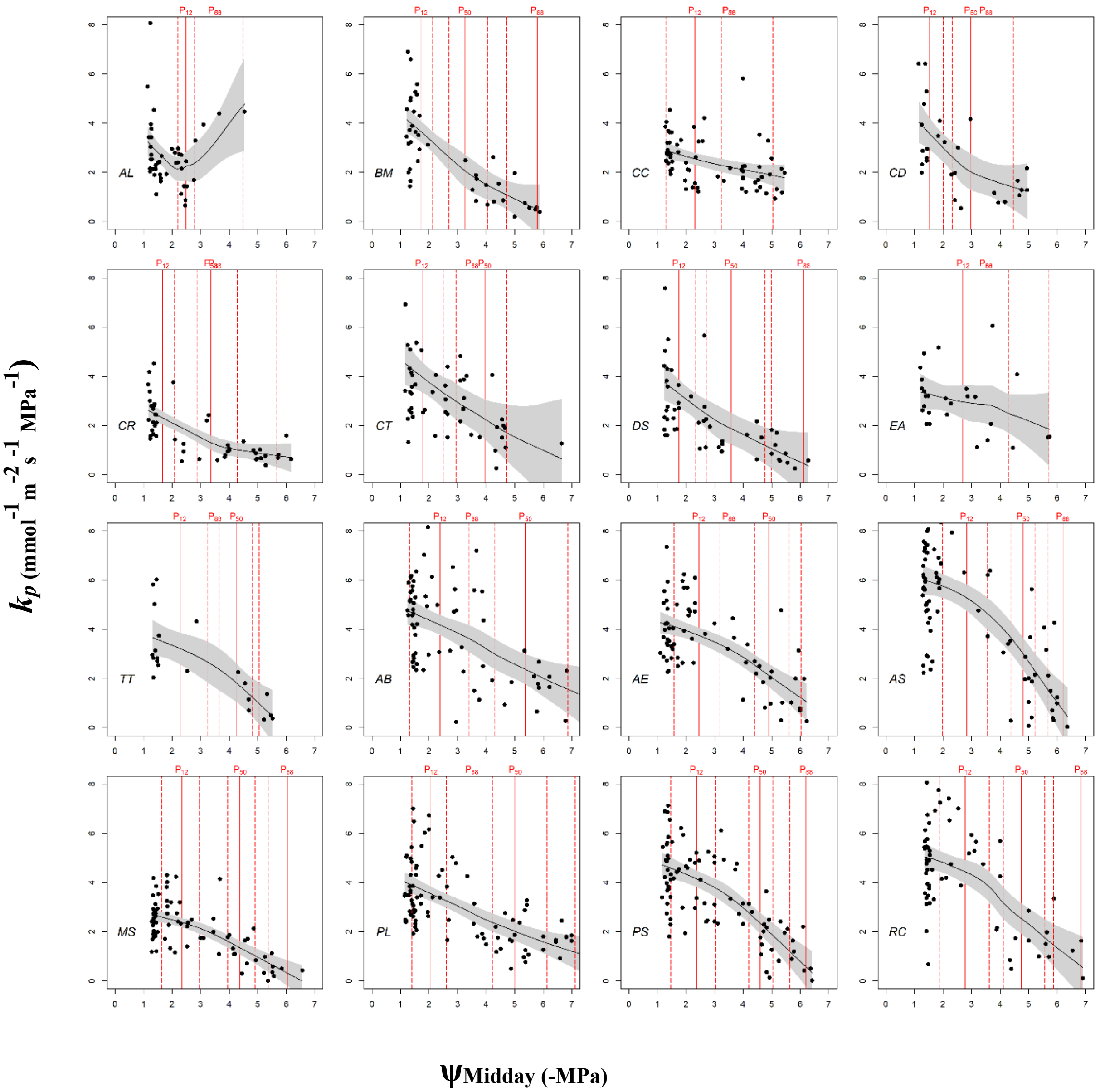


**Fig. S6** Loess fits for effective plant hydraulic conductance ($P_{kp12}$, $P_{kp50}$, $P_{kp88}$). For four species (AL, CC, CD and EA) $P_{kp88}$ values could not be fit. Red vertical lines show the fitted parameter value and confidence interval (CI) (2.5% and 97.5 %).

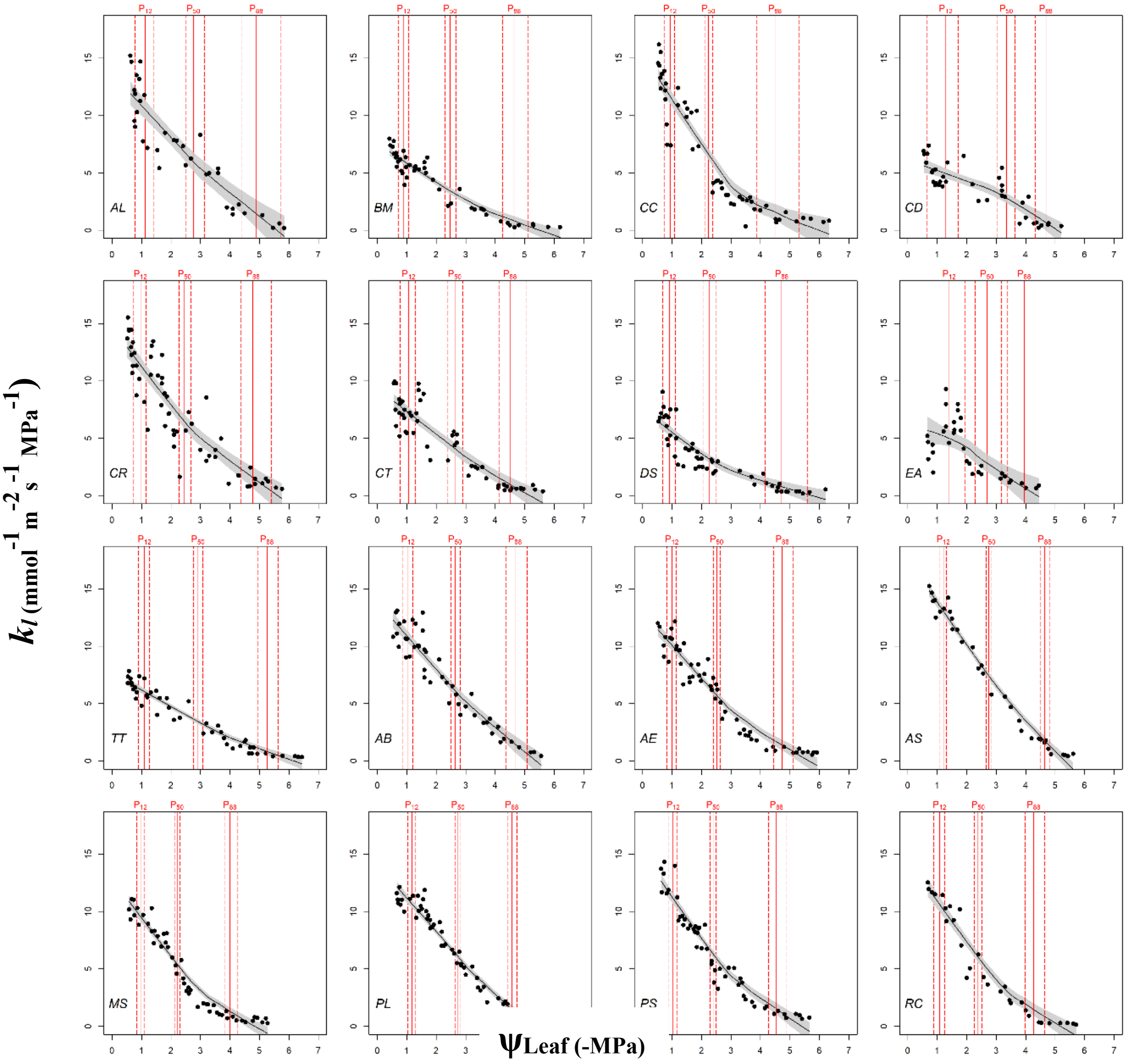


**Fig. S7** Loess fits for leaf hydraulic conductance ($P_{kl12}$, $P_{kl50}$, $P_{kl88}$). Red vertical lines show the fitted parameter value and confidence interval (CI) (2.5% and 97.5 %).

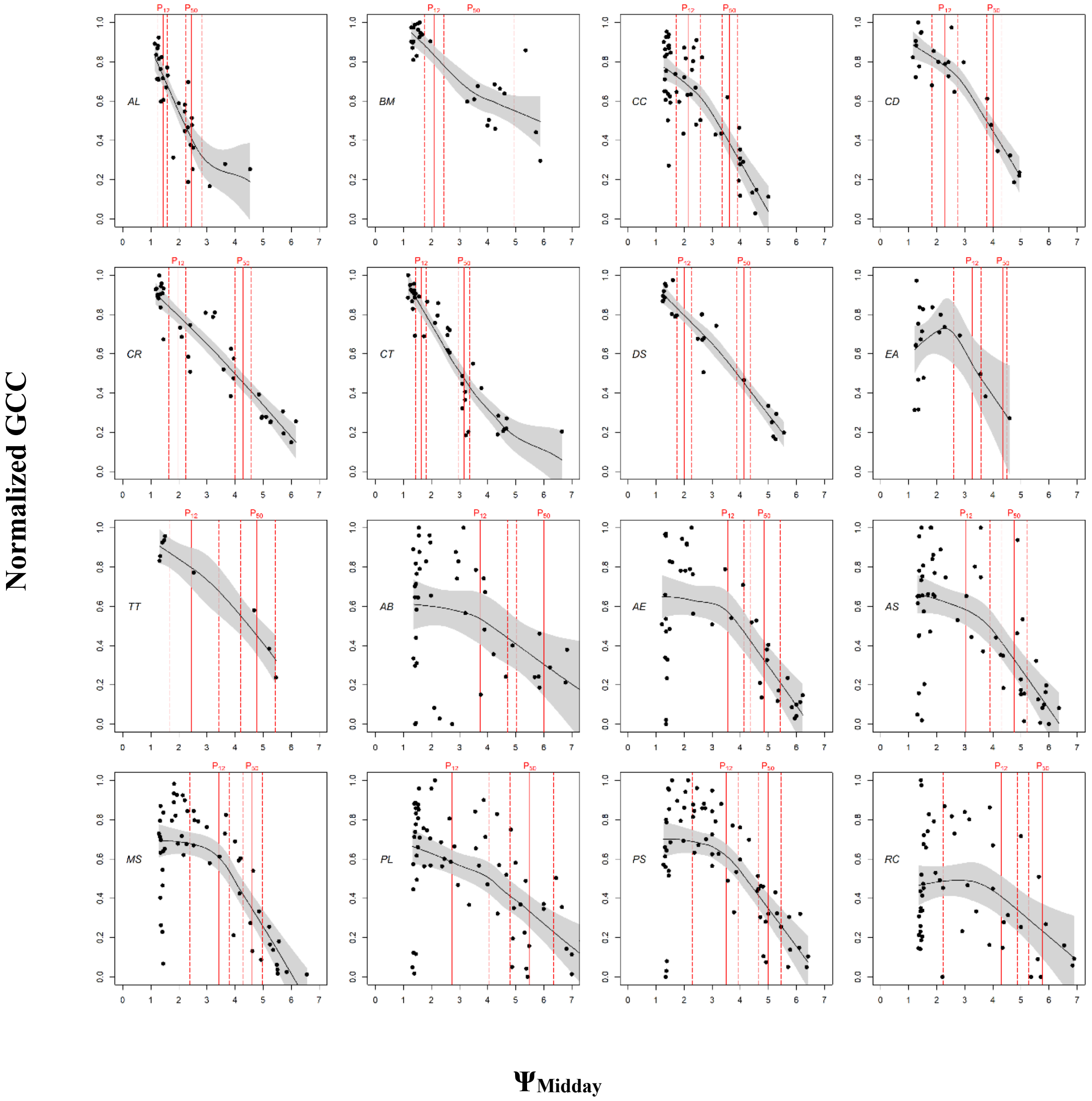


**Fig. S8** Loess fits for leaf browning using normalized GCC values. For each pixel of the images GCC was normalized to the range of zero to one. Note that the fits only retrieved $P_{Br12}$ and $P_{Br50}$ and not $P_{br88}$. Red vertical lines show the fitted parameter value and confidence interval (CI) (2.5% and 97.5 %).

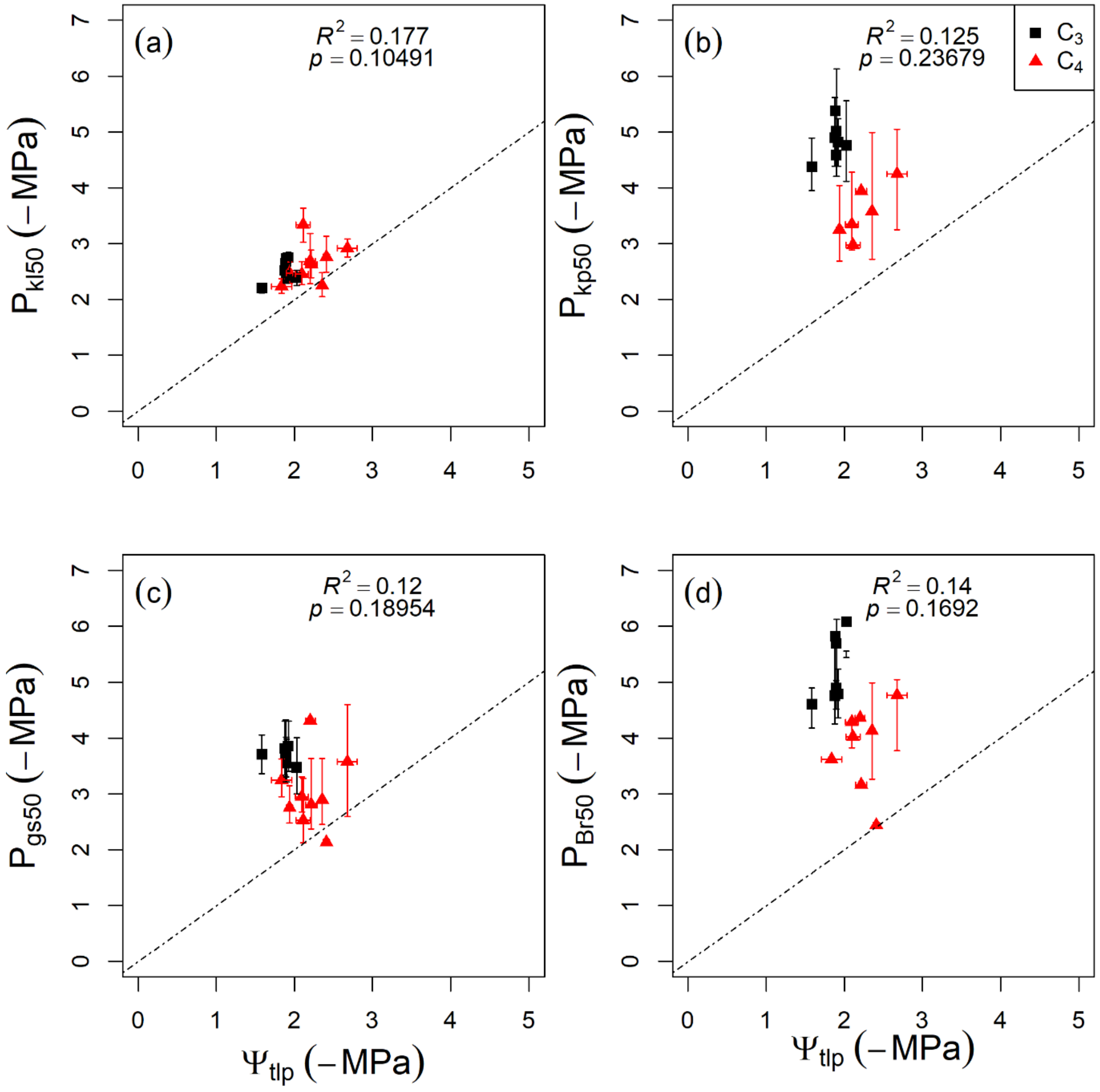


**Fig. S9** Relationships between turgor loss point ($\Psi_{tlp}$) and (a) leaf hydraulic impairment ($P_{kl50}$) (b) plant hydraulic impairment ($P_{kp50}$) (c) stomatal closure ($P_{gs50}$) traits and (d) leaf browning ($P_{Br50}$) traits.

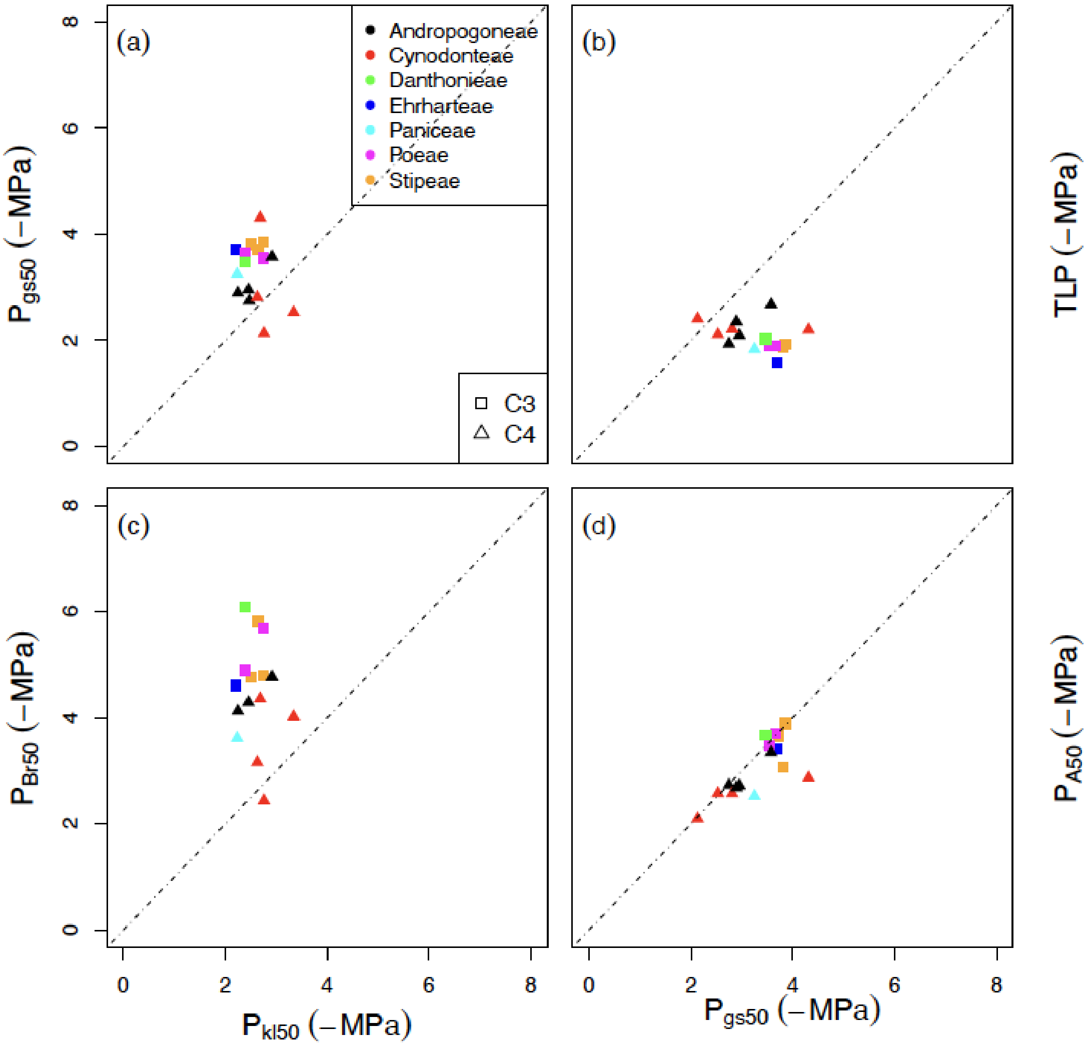


**Fig. S10** Same as Figure 4, but with points coloured to indicate grass tribe of each species.

**Table S1** Table of symbols, with units and their definitions

| Traits | Units | Definition |
|---|---|---|
| $\Psi_{leaf}$ | MPa | Leaf water potential |
| $g_s$ | mol $m^{-2}$ $s^{-1}$ | Stomatal conductance |
| $k_l$ | mmol $m^{-2}$ $s^{-1}$ $MPa^{-1}$ | Leaf hydraulic conductance |
| $k_p$ | mmol $m^{-2}$ $s^{-1}$ $MPa^{-1}$ | Plant hydraulic conductance |
| $C_{leaf}$ | mmol $MPa^{-1}$ $m^{-2}$ | Leaf capacitance |
| $A$ | μmol $m^{-2}$ $s^{-1}$ | Photosynthetic rate |
| $P_{kl12}$ | MPa | $\Psi_{leaf}$ at 12% loss of leaf hydraulic conductance |
| $P_{kl50}$ | MPa | $\Psi_{leaf}$ at 50% loss of leaf hydraulic conductance |
| $P_{kl88}$ | MPa | $\Psi_{leaf}$ at 88% loss of leaf hydraulic conductance |
| $P_{kp12}$ | MPa | $\Psi_{leaf}$ at 12% loss of leaf specific plant hydraulic conductance |
| $P_{kp50}$ | MPa | $\Psi_{leaf}$ at 50% loss of leaf specific plant hydraulic conductance |
| $P_{gs12}$ | MPa | $\Psi_{leaf}$ at 12% loss of stomatal conductance |
| $P_{gs50}$ | MPa | $\Psi_{leaf}$ at 50% loss of stomatal conductance |
| $P_{gs88}$ | MPa | $\Psi_{leaf}$ at 88% loss of stomatal conductance / stomatal closure |
| $P_{Br12}$ | MPa | $\Psi_{leaf}$ at 12% reduction of canopy/ leaf greenness |
| $P_{Br50}$ | MPa | $\Psi_{leaf}$ at 50% reduction of canopy/leaf greenness |
| $P_{A12}$ | MPa | $\Psi_{leaf}$ at 12% reduction of photosynthetic rate |
| $P_{A50}$ | MPa | $\Psi_{leaf}$ at 50% reduction of photosynthetic rate |
| $P_{A88}$ | MPa | $\Psi_{leaf}$ at 88% reduction of photosynthetic rate |
| TLP | MPa | $\Psi_{leaf}$ at leaf turgor loss – Turgor loss point |
| HSM | MPa | Hydraulic safety margin, calculated as the difference between leaf water potential at stomatal closure ($P_{gs88}$) and 50% loss of leaf hydraulic conductance ($P_{kl50}$) |
| $k_{lmax}$ | mmol $m^{-2}$ $s^{-1}$ $MPa^{-1}$ | Leaf maximum hydraulic conductance |
| $g_{smax}$ | mol $m^{-2}$ $s^{-1}$ | Maximum stomatal conductance |
| $k_{pmax}$ | mmol $m^{-2}$ $s^{-1}$ $MPa^{-1}$ | Plant maximum hydraulic conductance |
| $A_{max}$ | μmol $m^{-2}$ $s^{-1}$ | Maximum photosynthetic rate |
